# Dispersal of influenza virus populations within the respiratory tract shapes their evolutionary potential

**DOI:** 10.1101/2024.08.31.610648

**Authors:** Lucas M. Ferreri, Brittany Seibert, C. Joaquin Caceres, Kayle Patatanian, Katie E. Holmes, L. Claire Gay, Flavio Cargnin Faccin, Matias Cardenas, Silvia Carnaccini, Nishit Shetty, Daniela Rajao, Katia Koelle, Linsey C. Marr, Daniel R. Perez, Anice C. Lowen

## Abstract

Viral infections are characterized by dispersal from an initial site to secondary locations within the host. How the resultant spatial heterogeneity shapes within-host genetic diversity and viral evolutionary pathways is poorly understood. Here we show that dispersal within and between the nasal cavity and trachea maintains diversity and is therefore conducive to adaptive evolution, whereas dispersal to the lungs is stochastic and gives rise to population heterogeneity. We infected ferrets either intranasally or by aerosol with a barcoded influenza A/California/07/2009 (H1N1) virus. At 1, 2 or 4 days post infection, dispersal was assessed by collecting 52 samples from throughout the respiratory tract of each animal. Irrespective of inoculation route, barcode compositions across the nasal turbinates and trachea were similar and highly diverse, revealing little constraint on the establishment of infection in the nasal cavity and descent through the trachea. By comparison, infection of the lungs produced genetically distinct viral populations. Lung populations were pauci-clonal, suggesting that each seeded location received relatively few viral genotypes. While aerosol inoculation gave distinct populations at every lung site sampled, within-host dispersal after intranasal inoculation produced larger patches, indicative of local expansion following seeding of the lungs. Throughout the respiratory tract, barcode diversity declined over time, but new diversity was generated through mutation. De novo variants were often unique to a given location, indicating that localized replication following dispersal resulted in population divergence. In summary, dispersal within the respiratory tract operates differently between regions and contributes to the potential for viral evolution to proceed independently in multiple within-host subpopulations.

## Introduction

Dispersal from the initial site of infection to new susceptible locations is integral to viral propagation within infected hosts ^1,2^. Such expansion can yield a heterogenous population landscape, modulated by the number of viruses moving outwards from the original population, the distance traveled, and the frequency of dispersal events ^3–6^. These characteristics in turn shape the genetic composition of spatially separate sub-populations and thereby define the potential for selective and stochastic evolutionary processes to act ^7,8^. The generation of de novo mutations further increases potential for genetic divergence across the within-host landscape ^9–11^. Thus, the early stages of acute infection, during which viruses undergo extensive population expansion, may be highly consequential for viral evolutionary potential.

In ecological terms, expanding populations can exhibit a range of different dynamics. Diffusive expansions create a leading edge of individuals that move towards adjacent areas in contact with the source population ^12^. This mode of dispersal maintains genetic homogeneity, with smooth gradients of population density and genetic composition ^13,14^. Alternatively, punctuated expansion through long-distance dispersal events can yield isolated populations that spread locally ^15^. Under this dispersal regime, local spread is promoted by the availability of resources and lack of competition in empty habitats ^16^.

The dynamics of a population’s expansion directly impacts its evolution. Genetic drift and selection are notably sensitive to population size, since stochastic changes are more likely in small populations and deterministic processes are more efficient in large populations ^17^.

Following this principle, diffusive dispersal that maintains relatively large populations at the leading-edge should decrease the strength of genetic drift and allow for efficient selection. On the contrary, long-distance dispersal that generates small and isolated populations should lead to viral evolution being more strongly governed by genetic drift, that is, stochastic processes^14,18,19^.

In the case of respiratory viruses, infection is typically initiated by a small population and then drives a process of rapid spatial expansion throughout the respiratory tract ^3,20^. To understand how within-host dispersal from a focal population shapes opportunities for viral evolution, we monitored the spatial and temporal dynamics of a barcoded influenza A virus population during infection of ferrets. Our results link the population and evolutionary dynamics playing out within infected hosts and reveal extensive potential for differentiation of viral subpopulations during expansion within the respiratory tract.

## Results

### Extensive sampling allows mapping of virus dispersal across the respiratory tract

Potential for dispersal is high when the initial population is localized; conversely, potential is reduced when the initial population is distributed broadly such that the availability of uninfected areas is limited. To evaluate the evolutionary consequences of extensive dispersal during population expansion, we infected twelve ferrets either intranasally (giving a localized initial population) or by aerosol inhalation (giving a broadly distributed initial population). For intranasal inoculation, localized delivery was ensured by administering a small volume of 100 µL. The amount of virus delivered via the two routes was the same. To capture changes in dispersal dynamics over time, samples were collected at 1, 2, and 4 days post inoculation (dpi). To attain relatively high spatial resolution, extensive sampling of tissues throughout the respiratory tract was performed (Figure 1A). The nasal turbinates and trachea were collected in their entirety, dividing the organs into two and five sections, respectively. From the lungs, 45 sections were sampled across the six lobes: eight sections from each of the right cranial, right middle, right caudal, left caudal and left cranial and five sections from the accessory lobe.

**Figure 1.**
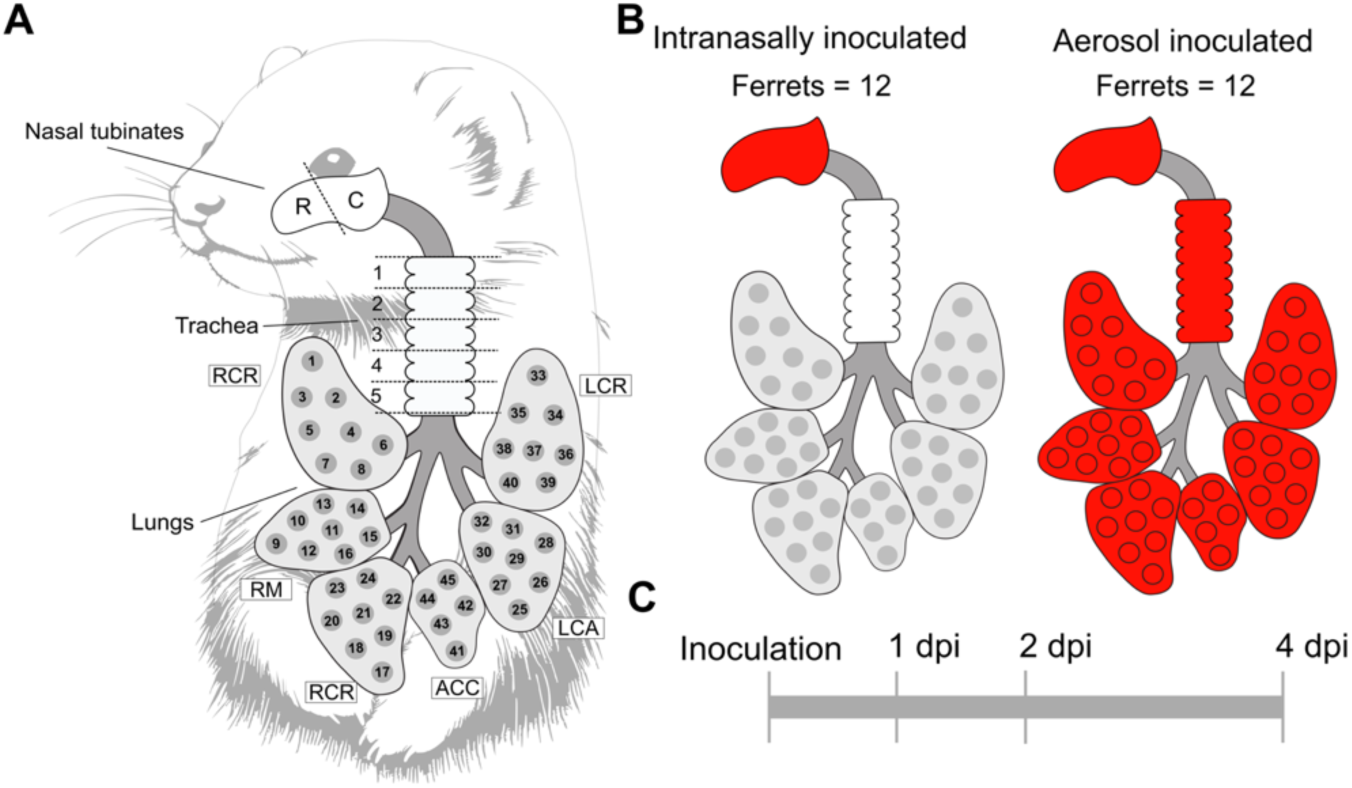
Experimental design for capturing spatial distribution of virus populations. A. Extensive sampling of infected ferrets included division of nasal turbinates into rostral (R) and caudal (C) sections, division of trachea into five sections (1-5) and collection of 45 sections across the six lung lobes: right cranial (RCR), right middle (RM), right caudal (RCA), accessory (ACC), left caudal (LCA) and left cranial (LCR). B. Two different inoculation techniques were used, intranasal and aerosol. C. Samples were collected at 1, 2, and 4 days post infection (dpi).

### Dispersal following intranasal inoculation produces discrete patches of infection in the lungs that expand as infection progresses

To evaluate the dispersal of virus populations, we quantified infectious virus present in each tissue section (Figure 2). Infection was well distributed in the nasal turbinate and tracheal sections at 1, 2 and 4 dpi. In the lungs, sparse patches of infection were detected at 1 dpi. Notably, most foci at this time covered small areas with low viral load, reflecting the initial phase of the infection at these sites. At later time points, ferrets 469 and 7843 showed large areas of infection in a subset of lobes (Figure 2). By 4 dpi, infection was more widespread in the lung but still not uniform; lung lobes showed wide variation in viral titers, including some sections that were negative for infectious virus. These data indicate that dispersal from the nasal cavity to the lungs occurs irregularly, such that virus is seeded unevenly. Subsequent expansion of the seeded populations is robust, but viral populations remain spatially heterogenous following this expansion.

**Figure 2.**
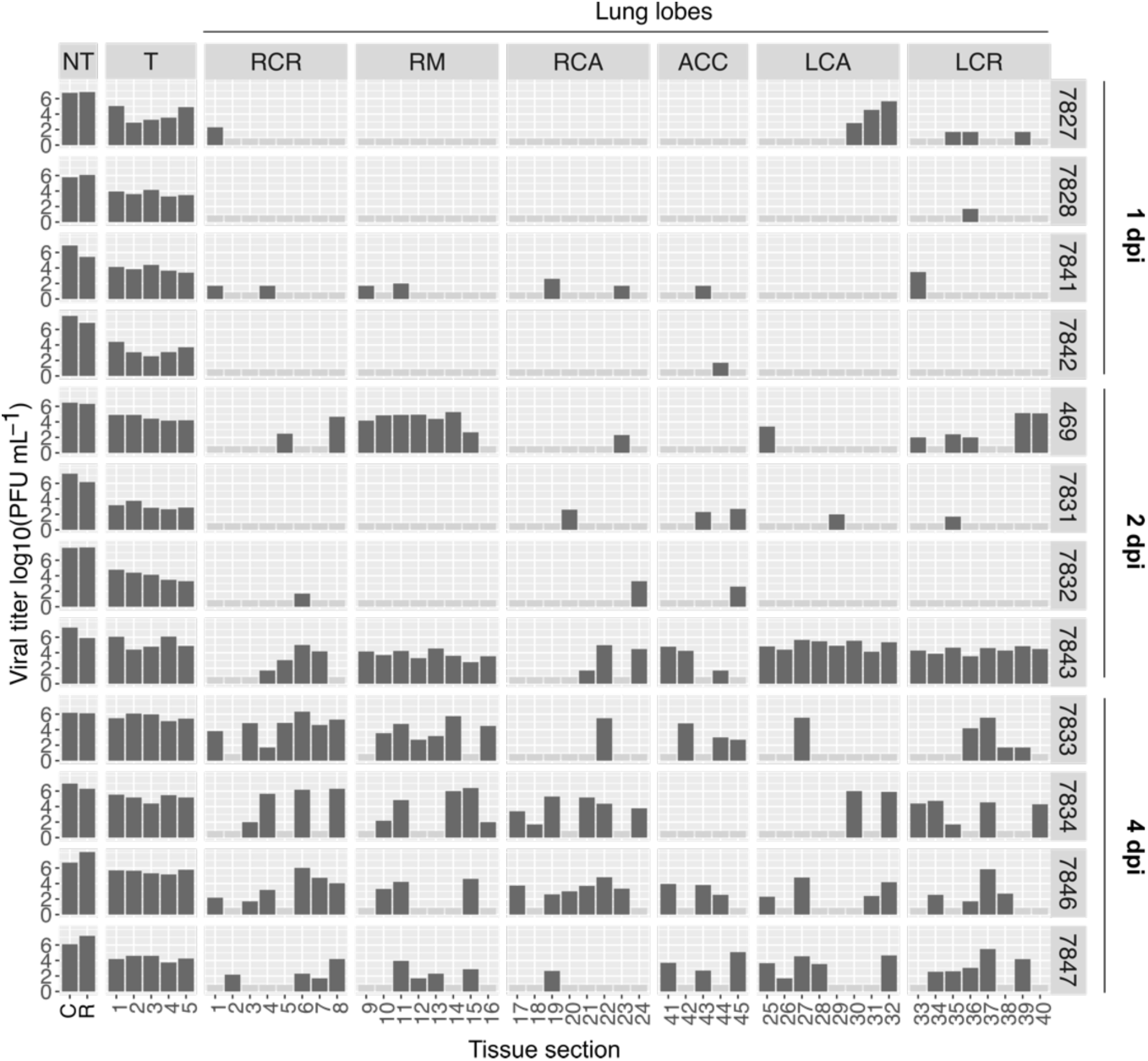
Distribution of viral populations in the lungs of intranasally inoculated ferrets. Dark grey bars depict positive samples. Eight sections from each lung lobe are shown, except the accessory lobe, which contains five sections. Lobes are designated as: RCR = right cranial; RM = right middle; RCA = right caudal; ACC = accessory; LCA = left caudal; LCR = left cranial. Numbers on the secondary y-axis show ferret identification numbers. Time of sampling is shown at the extreme right (dpi = days post-inoculation).

### Transfer of virus populations from the nasal cavity to the trachea is unconstrained

To evaluate how transfer from the source population influenced the genetic makeup of dispersing viral populations, we first compared the barcode composition present in the nasal turbinates and trachea of intranasally inoculated ferrets (Figure 3). In the nasal turbinates, populations were highly diverse at 1 dpi and showed modest differences between the rostral and caudal sections. In the trachea, populations were also highly diverse. Analysis of barcode species found in tracheas at 1 dpi indicated that the percentage of barcodes transferred from the nasal cavity to this organ was substantial: 90.1% for 7827, 47.6% for 7828, 41.1% for 7841, and 21.2% for 7842. In line with this observation, pairwise analysis of barcode compositions in nasal turbinate and tracheal sections revealed modest dissimilarity (Figure 3B). Looking within the trachea, consecutive sections showed similar barcode composition, suggesting that infection was seeded by the same population in a large portion of the organ. In contrast, pockets of low diversity suggestive of founder effects were also observed at the distal or proximal ends of the trachea in ferret 7827 (section 5), ferret 7828 (section 1) and ferret 7842 (section 5) (Figure 3A). These data suggest that the dispersal of virus from the nasal cavity to the trachea is typically unconstrained, such that stochastic changes in population composition during transfer to this anatomical are uncommon.

**Figure 3.**
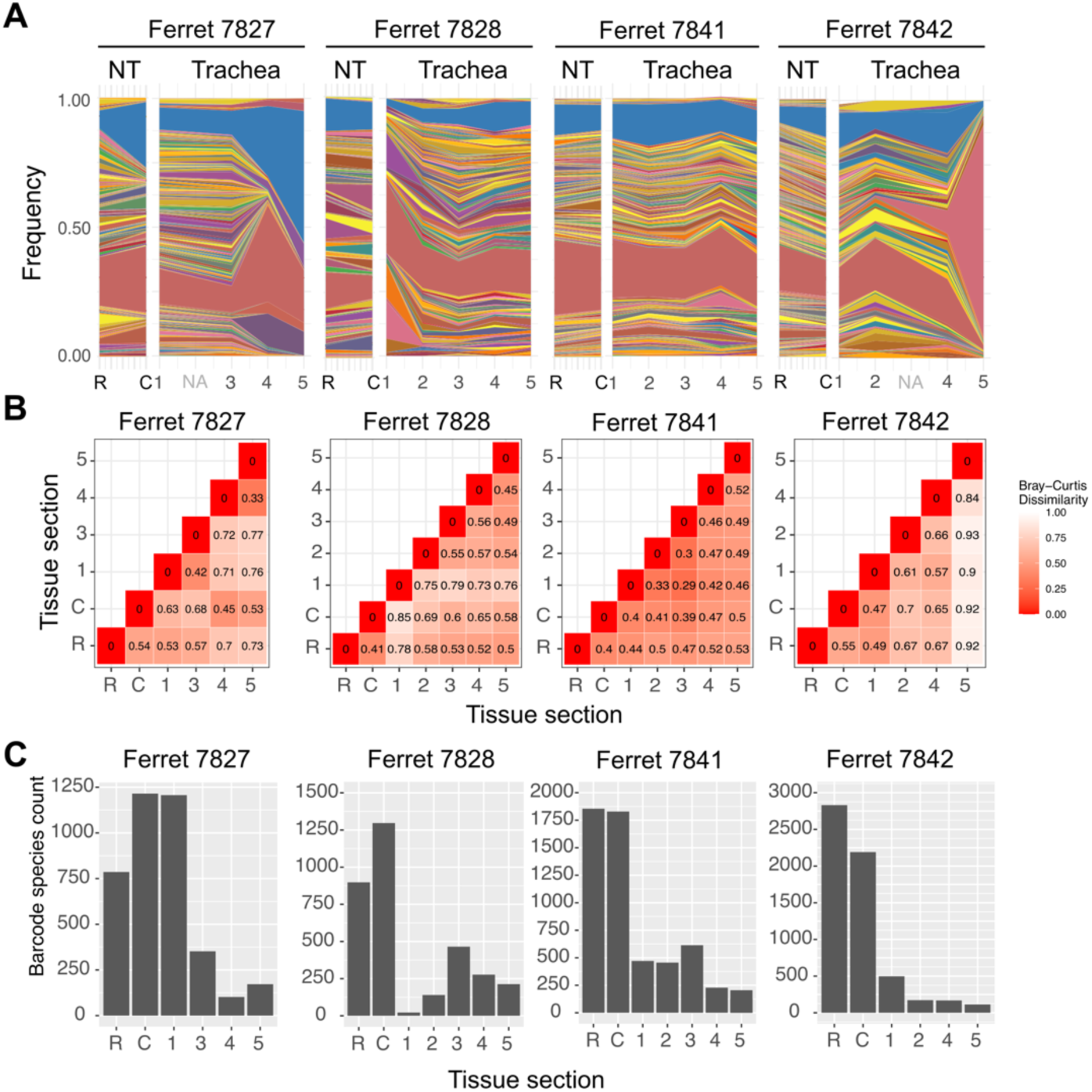
In intranasally inoculated ferrets, dispersal to the trachea is unconstrained and seeds a similar population throughout the organ. A. Barcode frequencies detected in the rostral (R) and caudal (C) sections of the nasal turbinates and sections 1 to 5 in the trachea (1-5) of four ferrets sampled at one day post-inoculation. Different colors represent unique barcodes. NA = not applicable; sequencing data are not available from these samples. B. Bray-Curtis dissimilarity index. Values within the matrices and the color scale show the extent of dissimilarity between viral populations sampled from different tissue locations. C. Number of unique barcode species detected in nasal turbinates and trachea.

### Transfer of viral populations to the lungs is accompanied by loss of diversity and establishment of genetically distinct subpopulations

To understand how dispersal over longer distances shapes the genetic composition of viral populations, we evaluated barcode composition in the lungs of intranasally inoculated animals (Figure 4 and Figure S2). Ferret 7843, at 2 dpi, displayed extensive infection in the lungs, providing an opportunity to evaluate the composition of viral populations both within and between lung lobes relatively early in the infection. At this early time, barcode composition is expected to maintain features of the seeding populations. In this ferret, all sections evaluated from lung lobes RM, LCA and LCR were positive for infection and each hosted a pauci-clonal viral population, indicative of a constriction in dispersing viral populations but not a tight bottleneck (Figure 4B). The distinct patterns of barcodes observed between lobes suggests that the RM, LCA and LCR lobes were seeded by different events, resulting in unique populations in each. To define the genetic relationships between viral populations occupying different sections within a lobe, we performed hierarchical clustering analysis (Figure 4C). The results show that the LCR lobe was infected by two different populations, the LCA lobe by three different populations and the RM lobe by two different populations (Figure 4C and D). The spatial proximity of genetically related populations further revealed that, once seeding occurred, populations went through extreme expansion, occupying large areas of the lung lobe (Figure 4F).

**Figure 4.**
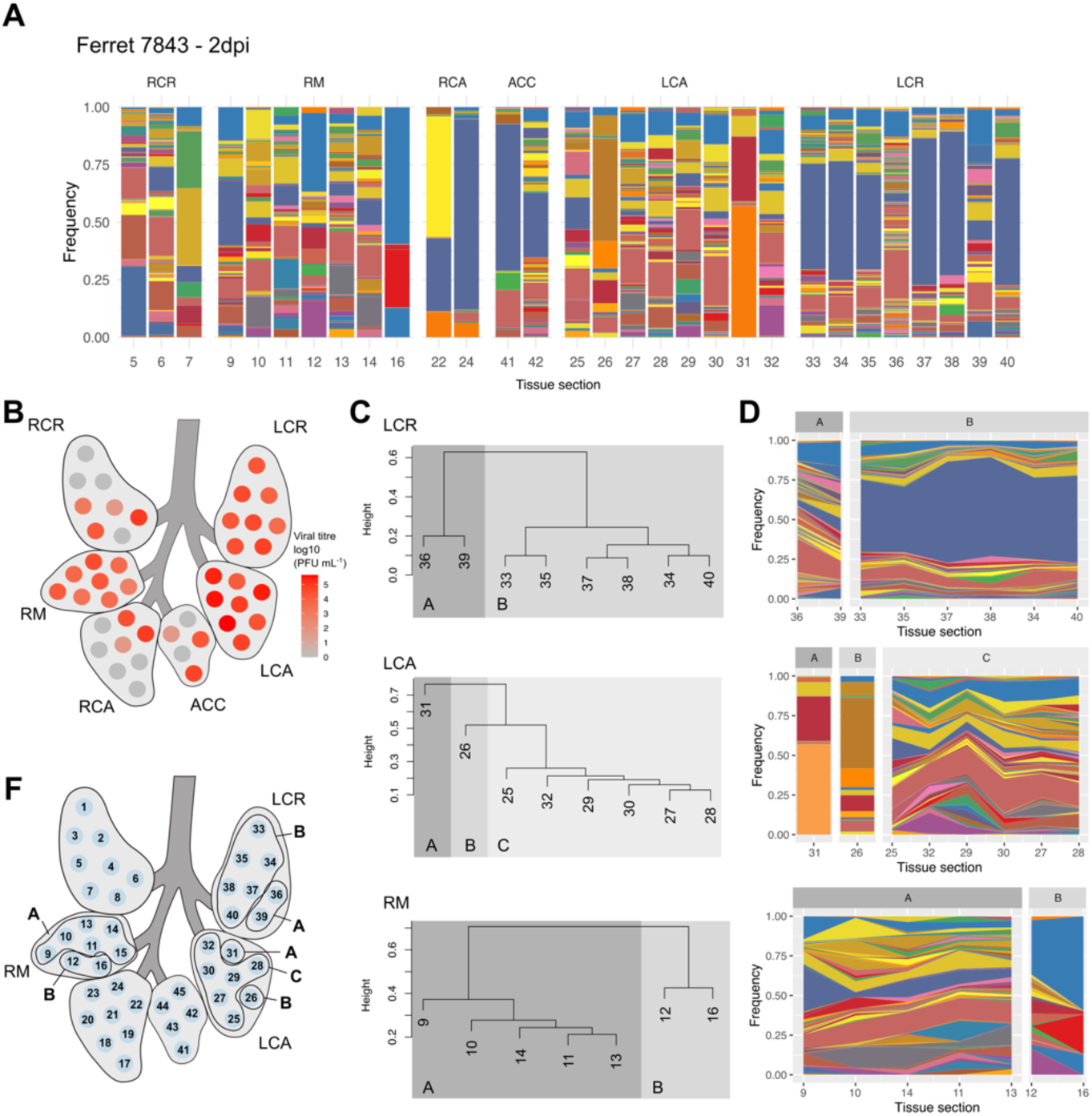
Expansion produces divergent populations in the lung lobes. All data shown are derived from intranasally inoculated ferret 7843, from which tissues were harvested at 2 days post inoculation. Related data from additional animals are found in Figure S2. (A) Distribution of barcodes within the lung. (B) Distribution of infectious virus in the lung. (C) Relationships among sampled viral populations as assessed by hierarchical clustering of barcode compositions. Height shows distance between clusters. (D) Continuity of barcodes within clusters identified in (C). (F) Spatial relationship of clusters within each lung lobe. RCR = right cranial; RM = right middle; RCA = right caudal; ACC = accessory; LCA = left caudal; LCR = left cranial.

### Standing diversity diminishes as infection progresses

Once infection is established, populations grow, influencing dispersal and the genetic makeup of the initial and new populations. We therefore investigated temporal trends in viral population dynamics by evaluating tissue samples collected at three different time points after intranasal inoculation. In both the nasal turbinate and tracheal tissues, barcode diversity significantly declined between 1 and 4 dpi (Figure 5A and B). This pattern was apparent at both sites despite differing trends in viral population size. Within the nasal cavity, population size remained constant throughout the experiment (Figure 5C), likely because the large initial population instilled at this site saturated the nasal cavity. Conversely, in the trachea, the population expanded throughout the experiment (Figure 5C). These results show that, independent of viral population size and growth, significant declines in barcode diversity occur over time in both the nasal cavity and the trachea. In lungs, a similar trend towards lower diversity at 4 dpi than 1 or 2 dpi was observed (Figure 5B and Figure S2). Overall, these data indicate that, as viral infection progresses, diversity declines.

**Figure 5.**
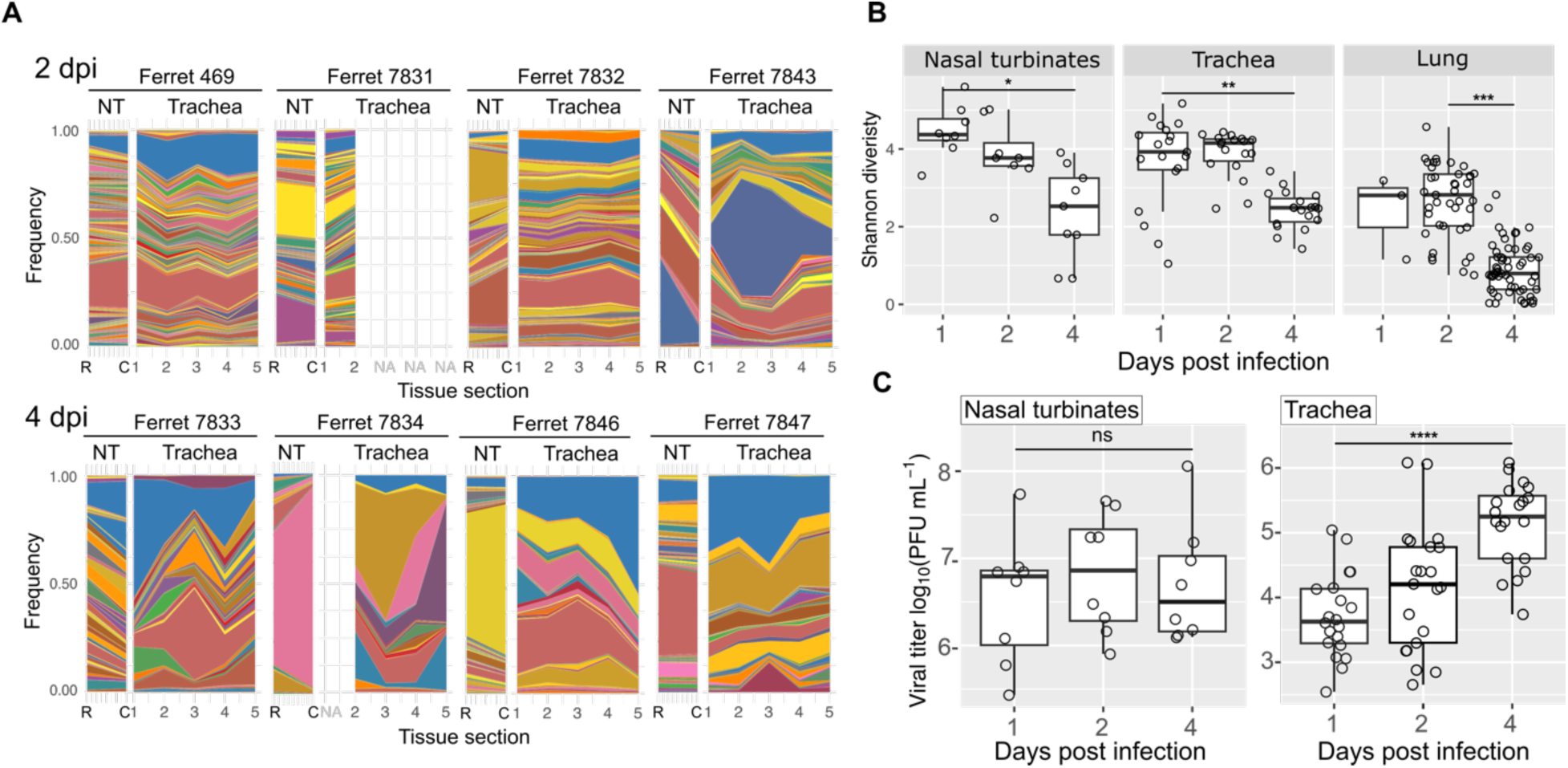
Barcode diversity declines over infection. (A) Barcode detection in nasal turbinates (NT) and trachea at 2 dpi and 4 dpi. (Data from 1 dpi are shown in Figure 3A.) (B) Shannon diversity in the nasal turbinates, trachea and lung. *P = 2.00e-03 comparing 1 and 4 dpi; **P = 3.66e-04 comparing 1 and 4 dpi; ***P = 4.77e-15 (comparing 2 and 4 dpi, due to the paucity of positive samples at 1 dpi). NA = not applicable. (C) Viral titers in nasal turbinate and tracheal sections. “R” = rostral section, “C” = caudal section. In the boxplots, the central line represents the median, the box edges represent the first and third quartiles (Q1 and Q3), and the whiskers extend to the smallest and largest values within 1.5 times the interquartile range (IQR) from Q1 and Q3. ****P = 7.77e-08.

### Heterogeneous populations arise following de novo diversification

As infections are established, error-prone replication of the viral genome produces new variants that can diversify populations. To evaluate whether in situ diversification led to the formation of spatially distinct subpopulations in intranasally inoculated animals, we sequenced whole viral genomes within the tissue samples collected. We then assessed the extent to which minor variants were shared between anatomical sites across the respiratory tract of the ferrets. We focused the analysis on the HA, NP, NA, M and NS gene segments due to their high coverage compared to the polymerase genes segments, increasing the confidence in the detected variants. Across multiple animals, minor variants emerged that were not detected elsewhere within the respiratory tract (Figure 6A and B). The extent of relatedness between populations varied, however. While the tracheal sections of some animals had high dissimilarity (e.g. ferret 7841), others shared some variants across sections, lowering the dissimilarity (e.g. ferret 7843 section 4 and 7847 sections 1, 2 and 3) (Figure S3). In the lungs, both pockets of high dissimilarity (e.g. section 31 in ferret 7843) and clusters of low dissimilarity (e.g. sections 34 and 35 of ferret 7843) were detected. Thus, the spatial distribution of variants formed de novo revealed in-tissue diversification that often produced localized subpopulations throughout the respiratory tract. Although the vast majority of variants outside of the barcode library were not detected in the instilled viral population, one prominent variant was carried over from the inoculum: HA K119N (Figure S4). The affected amino acid is localized in the head of the HA, outside of the receptor binding site. This variant was found in nasal turbinates and trachea at 1 and 2 dpi at frequencies similar to that observed in the inoculum, indicating that the processes of inoculation and passage of virus from the nasal turbinates to the trachea did not restrict the transfer of HA K119N. By 4 dpi, HA K119N was mostly detected above consensus frequencies in the tracheas. In the lungs, the frequency of HA K119N was variable without obvious trends. These data are consistent with the efficient transfer of barcode diversity from upper to lower respiratory tract and suggest that HA K119N may confer a minor selective advantage. The evolutionary dynamics of this variant appeared to be driven by positive selection in the trachea but by stochastic processes in the lungs.

**Figure 6.**
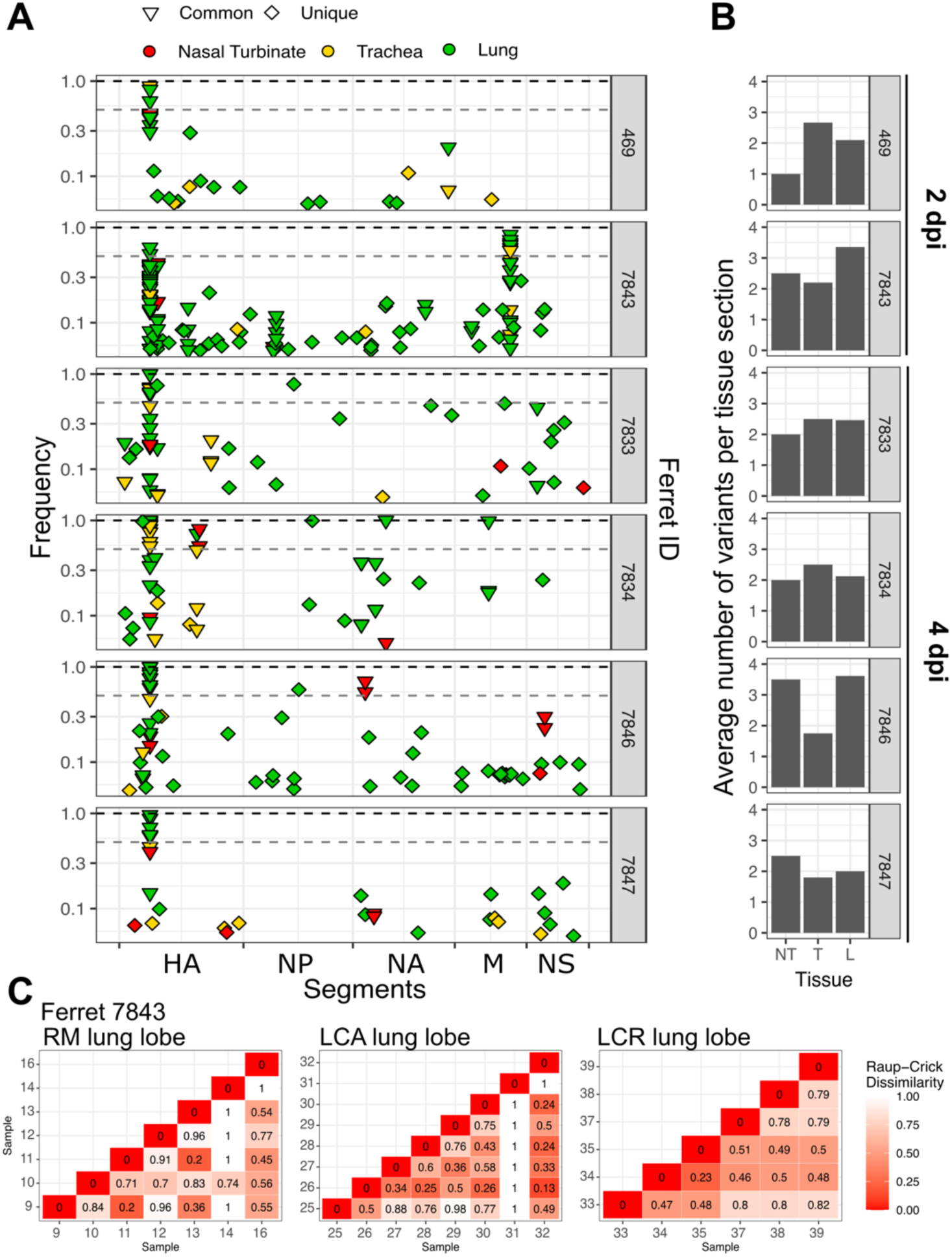
De novo mutations differentiate populations within tissues. (A) Minor variants detected across the respiratory tract of intranasally inoculated ferrets presenting extensive lung infection. Those designated as Unique (diamonds) were observed in only one tissue section. Those labelled as Common (inverted triangles) were observed in at least two sections. Variants comprising the introduced barcodes are not displayed. (B) Average number of variants per tissue section. (C) Raup-Crick dissimilarity index comparing lung sections in ferret 7843. Raup-Crick dissimilarity index was calculated without considering the barcode region.

### Aerosol inoculation results in rapid viral establishment within the lung, yielding isolated populations in the lower respiratory tract

When inoculum is deposited locally in the nasal cavity, infection of the lower respiratory tract arises through dispersal of virus over relatively long-distances, likely carried within particles of airway surface liquid that are small enough to penetrate the bronchial tree. By contrast, inoculation with aerosolized virus directly delivers virus throughout the respiratory tract, bypassing the process of within-host dispersal. We hypothesized that virus populations in ferrets infected via aerosols would show distinct spatial patterns from those infected intranasally, and that differences within the lower respiratory tract would reveal the implications of within-host viral dispersal for population genetic structure. Thus, ferrets were infected using infectious respiratory particles of 2.5 – 4 μm diameter, at the same dose delivered intranasally. Tissue samples were analyzed as before to define viral population size, spatial distribution and genetic composition.

Following aerosol delivery, high viral titers were observed across the respiratory tract already at 1 dpi. In stark contrast to animals inoculated intranasally, all four aerosol-inoculated ferrets sampled at this early time point exhibited extensive infection in each of the six lung lobes (Figure 7 A and C and Figure S5). Viral titers then declined at later time points (Figure 7 C and Figure S5).

**Figure 7.**
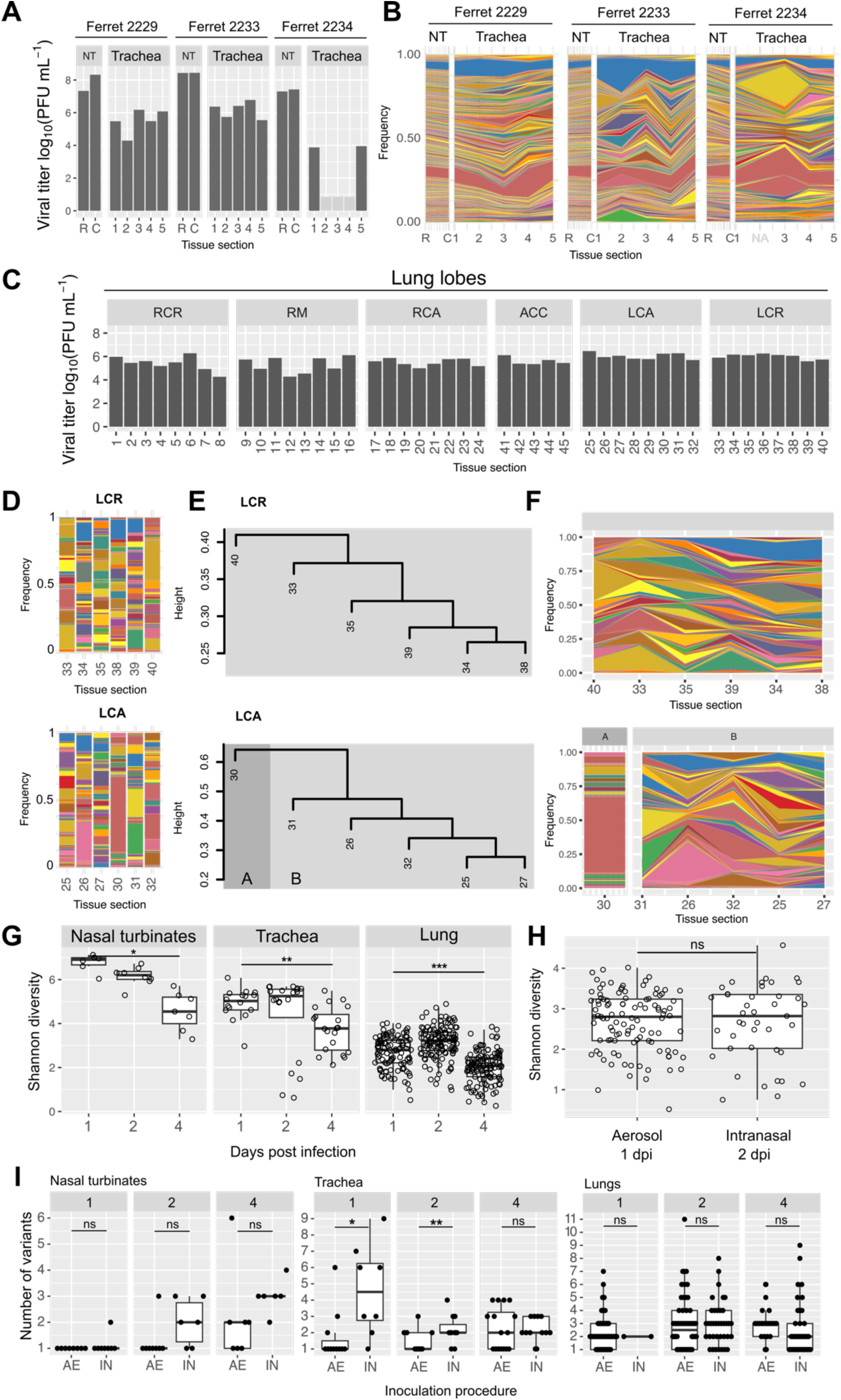
Dispersed viral delivery through aerosols seeds diverse, unique populations. (A) Virus titers in the nasal turbinates (NT) and tracheas of ferrets 2229, 2233 and 2234, sampled at 1 dpi. (B) Barcode detection in the NT and tracheas of ferrets 2229, 2233 and 2234. (C) Virus titers in the lung lobes of ferret 2230, sampled at 1 dpi. (D) Barcode composition by tissue section for the lung lobes LCA and LCR of ferret 2230. (E) Distance matrices of barcode frequencies were used to evaluate genetic relationships by hierarchical clustering. (F) Stacked plots organized according to the genetic relationships identified by hierarchical clustering for left caudal (LCA) and left cranial (LCR) lobes of ferret 2230. Related data (A-F) from additional animals are found in Figures S5-S7. (G) Shannon diversity in nasal turbinates, trachea and lungs. Comparison between 1 and 4 dpi: *P = 7.56e-4; **P = 7.02e-4; ***P = 6.04e-11. (H) Shannon diversity comparison between 1 dpi aerosol inoculated ferrets and 2 dpi intranasally inoculated ferrets. In the boxplots, the central line represents the median, the box edges represent the first and third quartiles (Q1 and Q3), and the whiskers extend to the smallest and largest values within 1.5 times the interquartile range (IQR) from Q1 and Q3. (I) Comparison of number of variants detected in ferrets inoculated intranasally (IN) or via aerosols (AE). *P = 0.023; **P = 0.030. Number in facets show days post inoculation.

Barcode analysis in nasal turbinate and tracheal sections showed high diversity at 1 dpi and lower diversity by 4 dpi, similar to patterns in intranasally inoculated ferrets (Figure 7G). In the lungs, the diversity established early was lower than that seen in nasal and tracheal tissues (P=1.19e-7, and P=5.58e-10 respectively). Of note, barcode diversity at each site in the lungs was comparable between aerosol and intranasally inoculated animals (Figure 7H). Each population sampled from aerosol inoculated lungs was however distinct, such that hierarchical clustering based on barcode composition revealed few clusters with heightened relatedness. Further evidence of spatial heterogeneity was apparent from the rugged intersections between tissue sections in stacked plots showing barcode compositions (e.g. in the RM, LCR and LCA lung lobes of ferret 2230) (Figure 7D-F and Figure S6). Thus, inoculation with aerosolized virus established a robust, distributed infection in the lungs that comprised many highly localized subpopulations. As seen with dispersal from the nasal cavity, these subpopulations were pauci-clonal, revealing a moderate loss of diversity during penetration of virus into the lungs. In contrast to lung subpopulations within intranasally inoculated animals, however, virus that established in lungs following aerosol inoculation did not spread to contiguous sites. These observations suggest that aerosol-seeded subpopulations had little opportunity to disperse to new susceptible locations due to the synchronized delivery of virus throughout the respiratory tract.

## Discussion

Using both genetic barcodes and de novo mutation, we examined the extent to which influenza virus dispersal within the respiratory tract leads to genetic divergence of spatially separated populations. Incorporation of barcodes allowed detection of changes in population composition during the establishment, propagation and clearance of infection, while analysis of whole genome sequences allowed the spatial-temporal mapping of viral diversity generated during infection ^21–25^. The use of two contrasting inoculation methods furthermore allowed the impact of initial seeding location to be investigated. The distribution of viral genotypes throughout the respiratory tract and over time revealed a complex landscape, with stochastic changes in viral population composition dominant in the lungs and conditions more conducive to selective evolution prevailing in the nasal cavity and trachea.

Within the nasal cavity, high barcode diversity with modest differences between sections was observed, signaling efficient establishment of virus populations by both delivery methods. Viral populations in the trachea at 1 dpi were large and showed high barcode diversity. Seeding was furthermore likely to have occurred simultaneously across the trachea using both inoculation routes, as sections shared highly similar barcode composition. The similarity of patterns in the trachea between intranasal and aerosol inoculation groups demonstrates the efficiency of dispersal from nasal cavity to the trachea, showing a lack of genetic bottleneck. Indeed, these data suggest that, when infection initiates in the nasal cavity, the trachea is populated via diffusive dispersal ^13^. Dispersal by diffusion occurs when the leading edge of expansion remains continuous with the original population, resulting in modest genetic differences between these areas ^12^. This form of dispersal can lower the impact of genetic drift by establishing a large initial population: founder populations with high diversity and rapid growth are likely to undergo selective processes ^26,27^. Thus, the diffusive dispersal and rapid growth of virus populations in the trachea suggests that this anatomical location can act as a site for viral adaptation.

Prior work examining the population genetic structure of influenza virus in the lower respiratory tract revealed spatial heterogeneity at the level of individual lung lobes ^28,29^. However, the structure of populations on a smaller spatial scale has not been examined, leaving the number of seeded populations per lung lobe and the features of their local expansion uncharacterized. We saw that, following intranasal inoculation, dispersal to the lungs initially resulted in sparse and small foci. Some lobes were more rapidly colonized than others, an effect that appeared to be stochastic since trends were not apparent across animals. These features are characteristic of long-distance dispersal events in which few individuals establish a new population far from the origin ^30^. Once lungs were seeded, we observed that populations increased in density and expanded spatially. In some cases, contiguous tissue sections showed similar genetic composition, revealing diffusive dispersal within a lung lobe. By contrast, some sections were genetically distinct from adjacent sites and, when infection was seeded through aerosol inoculation, all lung sections were distinct from adjacent sites. The genetic structure detected within the lung lobes suggests that founder populations were relatively small and multiple independent seeding events were followed by little to no mixing between foci. Such nonhomogeneous dispersal is conducive to genetic drift ^19^ and can rapidly lead to population differentiation at small spatial scales ^31^. In the infected respiratory tract, dispersal may be heterogeneous as a result of the structure of the lung ^32^, super infection exclusion ^33^ and/or innate immune activation ^34^.

Throughout the respiratory tract and following both modes of inoculation, substantial de novo variation was detected. When considered on a per-sample basis, the extent of variation did not differ between inoculation modes or across nasal turbinates, trachea and lungs, indicating that the force of purifying selection was comparable. This observation is perhaps surprising given the above-noted differences in dispersal: long distance dispersal has been proposed to weaken purifying selection due to high rates of genetic drift at the leading edge (Bosshard et al. 2017). We hypothesize that the similarity of variant accumulation across sites is the result of the relative timing of dispersal and variant emergence: dissimilarity in variant profiles across tissue sites suggests that variants accumulated largely following long-distance dispersal. Furthermore, barcode composition suggests that early long-distance dispersal to the lungs was followed by diffusive within-lobe expansion, similar to that in nasal and tracheal sites.

Certain limitations in our experimental design should be considered. First, how the two modes of inoculation analyzed relate to natural infection is unclear. While a range of modes are likely to occur in nature, the spatial distribution of seeding viral populations shapes subsequent spatial dynamics. Second, we used ferrets, which present both similarities and differences to humans and other natural hosts of influenza A virus ^35,36^. The ferret’s relatively long trachea, quadrupedal posture and immunological naivety are just a few of its features that may shape viral dispersal dynamics. Third, while many samples were collected, these were too large to consider viral dynamics at a fine scale; sections may have captured multiple seeded populations. Finally, as with any analysis reliant on next generation sequencing, our ability to detect minor variants was limited ^9^. In this case, we applied a frequency threshold of 5% and excluded analysis of the polymerase gene segments due to low coverage. The measures were designed to limit artifacts but reduce our capacity to detect shared and unique variants.

In summary, our data reveal that diffusive dispersal within the nasal cavity and trachea maintains population diversity, whereas long distance dispersal to the lungs is associated with loss of diversity within seeded subpopulations. Once viral populations are established at a given site, there is little mixing among foci; consequently, both founder effects during long distance dispersal and subsequent diversification result in population heterogeneity. The distinct features of local viral populations in the upper and lower respiratory tracts suggest that different evolutionary forces are likely to prevail in each: while maintenance of diversity is conducive to selection, stochastic loss of diversity during seeding of the lungs will often impede selection. Thus, by defining population genetic structure, the spatial expansion of infection throughout the respiratory tract is expected to yield a complex and varied evolutionary landscape.

## Acknowledgments

We thank the Emory Integrated Genomics Core (EIGC), particularly Lyra Griffiths, and the Emory Primate Center Genomic Core (EPC), particularly Kathryn Pellegrini and Steven Bosinger. We also would like to thank Dr. David VanInsberghe for valuable discussions and suggestions. The research was funded by NIH R01 AI154894 (A.C.L., K.K., D.R.P.) and the NIAID Centers of Excellence for Influenza Research and Response (CEIRR), contract number 75N93021C00017 (A.C.L., K.K.) and contract number 75N93021C00014 and Options 15A, 15B and 17A (D.R.P). Additional funds were provided to D.R.P. by the Georgia Research Alliance and the Caswell S Eidson Chair in Poultry Medicine endowment funds. Additional support was provided by Flu Lab (N.S., L.C.M.).

## Author Contributions

L.M.F. contributed to the conception of work, experimental design, analysis, and interpretation of data, writing of manuscript drafts and editing; B.S. and J.C. contributed to experimental design and data acquisition. K.P. contributed to data acquisition. K.H. contributed to experimental design and data acquisition. C.G., F.C.F., S.C, contributed to data acquisition. N.S. and L.M. contributed to experimental design. K.K. contributed to funding acquisition and the conception of the work. D.R.P. contributed to funding acquisition, the conception of the work, experimental design, and interpretation. A.C.L. contributed to funding acquisition, the conception of the work, experimental design, data analysis, and interpretation, writing of manuscript drafts and editing.

## Materials and Methods

### Cells

A seed stock of MDCK cells at passage 23 was expanded and maintained in Minimal Essential Medium (Gibco) with 10% fetal bovine serum (FBS; Atlanta Biologicals) and Normocin (Invivogen). 293T cells (ATCC, CRL-3216) were maintained in Dulbecco’s Minimal Essential Medium (Gibco) with 10% FBS and Normocin. All cells were cultured at 37°C and 5% CO_2_ in a humidified incubator. The cell lines were not authenticated. While in use, cells were tested monthly for Mycoplasma contamination. The medium for culturing influenza A virus in MDCK cells (virus medium) was prepared by supplementing Minimal Essential Medium with 4.3% bovine serum albumin (BSA; Sigma) and Normocin.

### Viruses

The Cal/09 WT and Cal/09 NA-BC viruses were derived from influenza A/California/07/2009 (H1N1) virus and were generated by reverse genetics. Briefly, co-cultures of 293T and MDCK cells were seeded 16-24 hours prior to transfection. An eight plasmid reverse genetic system was used for virus rescue ^37^. Cells were transfected using XtremeGene 9 reagent (Sigma). After 40 h at 33°C, the recovered virus was propagated in MDCK cells to create working stocks. To avoid the accumulation of defective viral genomes, propagation was performed from a low MOI, ensuring a sufficient viral population size to maintain barcode diversity. Titration of both the stocks and experimental samples was conducted by plaque assay in MDCK cells. The diversity of the barcodes in the virus stock was confirmed through next-generation sequencing (Figure S1).

### Generation of Cal/09 NA-BC plasmid

The genomic region of Cal/09 virus into which the barcode was engineered was selected by aligning 64 sequences of H1N1-subtype influenza A viruses isolated between 2006 and 2014. A region with numerous nucleotide substitutions was identified in the NA segment from genomic position 449 to 509. Twelve synonymous mutations were identified within this region and the barcode was produced by incorporating this natural variation. A double-stranded Ultramer (IDT) was designed that contained degenerate bases with two possible nucleotides at each of the twelve chosen sites (5’-AA**R**CATTC**M**AATGG**R**ACCAT**W**AA**R**GACAG**R**AG**Y**CC**W**TATCG**R**AC**Y**CTAATGAGCTGTCC**Y** AT**W**G-3’).

Site-directed mutagenesis was employed to insert an XhoI restriction site within the barcode region of the wild-type reverse genetics plasmid, pHW Cal/09 NA, before barcode insertion. This step was necessary to allow restriction digestion of the parental template. Successful mutagenesis was confirmed through XhoI restriction digest. Furthermore, two stop codons were introduced in the barcode region to truncate the open reading frame, rendering the wild-type pHW Cal/09 NA nonfunctional. These modifications were placed to minimize wild-type carry-through in cloning and virus generation steps. Modifications were confirmed by Sanger sequencing.

To generate a linearized template for barcode insertion, the following steps were performed: XhoI digestion of the plasmid stock was followed by phosphatase treatment with rSAP (NEB) to dephosphorylate the cut ends of the plasmid. The plasmid was then amplified by PCR using primers that extend outward from the barcode region: Ca07_LINEAR_NA_511_F 5’-GTGAAGTTCCCTCTCCATACAACTCAAGATTTGAG-3’ and Ca07_LINEAR_NA_446_R 5’-GTCATTTAGCAAGGCCCCTTGAGTCAAG-3’. The linearized PCR product was isolated through PCR purification (QIAquick PCR Purification Kit, Qiagen), followed by a dual digestion with DpnI and XhoI to remove residual WT plasmid. Another round of PCR purification was conducted, followed by an assembly reaction using the NEBuilder HiFi DNA Assembly Kit (NEB) to insert the Ultramer into the linearized vector and re-circularize it.

The product was then transformed into DH5-α cells (NEB). After plating onto LB-amp plates, approximately 1 × 10^4 colonies were collected and pooled into LB-amp culture media, then incubated at 37°C for five hours before harvesting the bacterial population for plasmid purification by Maxiprep (Qiagen Plasmid Maxi Kit). The presence of a diverse barcode in the plasmid stock was verified by next-generation sequencing (Figure S1). This plasmid stock was then used to generate Cal/09 NA-BC virus by reverse genetics in combination with seven plasmids encoding Cal/09 WT gene segments in a pDP2002 vector.

### Ferrets

Ferret studies were approved and conducted in compliance with all the regulations stated by the Institutional Animal Care and Use Committee (IACUC) of the University of Georgia (AUP2022 04-026-A3). Studies were conducted under ABSL-2 conditions at Biological Sciences, University of Georgia. Animal studies and procedures were performed according to the Institutional Animal Care and Use Committee Guidebook of the Office of Laboratory Animal Welfare and PHS Policy on Humane Care and Use of Laboratory Animals. Animal studies were carried out in compliance with the ARRIVE guidelines (https://arriveguidelines.org). Twenty-week-old male and female ferrets were acquired from Triple F Farms (Gillett, PA, USA). One day after arrival, ferrets were anesthetized with an i.m. injection (0.5 ml/kg) of a ketamine/xylazine cocktail (20 mg/kg ketamine, 1 mg/kg xylazine) in the thighs, followed by blood collection to test serum samples prior to influenza A virus infection using NP ELISA (IDEXX, ME, USA). All ferrets tested negative for previous influenza A virus exposure. A subcutaneous implantable temperature transponder (BMDS, DE, USA) was inserted in the back of the neck to identify and monitor the body temperature of each ferret. Ferrets were acclimated for seven days upon arrival before virus inoculation.

### Infection of ferrets with barcoded Cal/09 NA-BC virus via intranasal inoculation

Ferrets were anesthetized as described above and inoculated intranasally with 1 x 10^6^ PFU/ferret in a total volume of 100 µL. Ferrets were monitored for clinical signs daily. On day 1-, 2-, and 4-days post-inoculation (dpi), a subset of four ferrets (two male and two female) were anesthetized and humanely euthanized using 1 ml of Euthasol® (Virbac, TX, USA). Necropsies were subsequently performed to collect two sections of nasal turbinates, five sections of the trachea, and forty-five sections of the lungs using 4 mm Integra™ Miltex™ Standard Biopsy Punches. Samples were frozen at −80°C until processed.

### Infection of ferrets with barcoded Cal/09 NA-BC virus via aerosol inoculation

For aerosol inoculation, the nose-only bioaerosol exposure system included a Buxco mass dosing controller (Data Sciences International, MN, USA), Aeroneb lab control module (Kent Scientific, CT, USA), Aeroneb lab nebulizer unit (Small VMD; Kent Scientific, CT, USA), stackable inhalation tower with seven ports (Data Sciences International, MN, USA), nose-only ferret restraints with ally neck restraints (Data Sciences International, MN, USA), HEPA-CAP filters (VWR, PA, USA) and multiples sizes of plastic tubing and tubing adapters. Before exposure, ferrets were anesthetized as described above. During each aerosol inoculation, two ferrets of the same sex were exposed to aerosolized virus inoculum simultaneously for 10 min, in which two ports were connected to the nose-only ferret restraints. Throughout the exposure, 5 L/min of HEPA-filtered air was provided by the Buxco mass dosing controller to maintain an input of air that was more than double the resting respiratory minute volume of both ferrets. The flow rate of inputted air was measured at the start of each experimental exposure. For aerosolization of virus inoculum, 4 mL of the virus diluted in 1X PBS was aerosolized using the Aeroneb lab nebulizer with an expected particle size of 2.5 −4 µm at a rate of 0.409 mL/min. Concurrently, exhaled air was removed through two outflow tubes connected to a HEPA filter. The concentration or titer of virus inoculum used in the nebulizer (*C_neb_* in units of PFU/mL) to achieve a desired final dose (*D* = 1 x 10^6^ PFU) was calculated according to Equations 1 and 2:

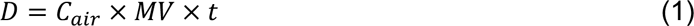

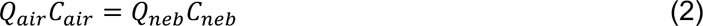

where *C_air_* is the virus concentration in air (PFU/L), *MV* is the respiratory minute volume estimated according to Equation 3, *t* is the exposure time of 10 min, *Q_air_* is the flow rate of air into the exposure system of 5 L/min, and *Q_neb_* is the flow rate of liquid through the nebulizer of 0.409 mL/min. Solving Equation 2 for *C_air_* and substituting this expression into Equation 1 produces an expression that can be rearranged to solve for *C_neb_* as a function of *D*, *MV*, *t*, *Q_air_*, and *Q_neb_*. This calculation assumes no inactivation of virus during nebulization and exposure.

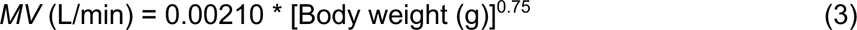

Due to the difference in weight among males and females, different volumes of inoculum were used for each sex, maintaining total infectious units per animal. Aerosol exposures were conducted at room temperature (20-22°C) and 50-60% relative humidity. After exposure, the nose of each ferret was wiped to limit the inoculation of alternate routes. On day 1-, 2-, and 4-days post-inoculation (dpi), a subset (n=2/sex/day) of aerosol-inoculated ferrets were anesthetized as previously described, bled, and humanely euthanized using 1 ml of Euthasol® (Virbac, TX, USA). Necropsies were subsequently performed to collect two sections of nasal turbinates, five sections of the trachea, and forty-five sections of the lungs. Samples were frozen at −80°C until processed.

### Tissue homogenization for RNA extraction

Stainless steel beads (5 mm, Qiagen, Cat no. 69989) and 2.0 ml bead-beating tubes (Olympus Plastics, Genesee Scientific, Cat. No. 21-253) were used for tissue homogenization. The tubes were pre-filled with PBS plus Normocin (InvivoGen) and one steel bead each. Homogenization was performed using the MP Fastprep 24 Homogenizer (MP, Cat. No. 116004500) set to 6 m/s for 60 seconds. The homogenized samples were then centrifuged at 15,000 RCF for 5 minutes at 4°C. The clarified homogenate was aliquoted and stored at −80°C.

### Sample processing for next generation sequencing of Cal/09 neuraminidase barcode region

RNA was extracted from a 145 μL volume of the clarified supernatant of each homogenized sample or virus stock using NucleoMag RNA isolation (Macherey-Nagel) either manually or in a EpiMotion 5075vt liquid handler. Next, PCR amplification targeting the barcode region was performed using 2.5 μL of purified RNA and One-Step Superscript III Platinum Taq Mix (Invitrogen). The thermocycler amplification protocol was: 55°C for 2 min, 45 °C for 60 min, 94 °C for 2 min, then 30 cycles of 94°C for 30 s, 59°C for 30 s, 68°C for 1 min and 68°C for 5 min. Primers: Cal07_NA_bc_eigc F 5’ **TCGTCGGCAGCGTCAGATGTGTATAAGAGACAG**ACCTTCTTCTTGACTCAAGGG 3’ and Cal07_NA_bc_eigc R 5’ **GTCTCGTGGGCTCGGAGATGTGTATAAGAGACAG**CAAGCGACTGACTCAAATCTTGA 3’.

In bold are the sequences pertaining to the Illumina adapters. Amplicons were purified using Agencourt AMPure XP magnetic beads with a 30 μL elution in water. Bead to sample volume ratio was 2.5. Clean PCR amplicons were sequenced at the Emory Integrated Genomics Core. Library preparation was performed with omission of the tagmentation step and sequencing was performed on a MiSeq (Illumina) platform. Amplicons were sequenced using the MiSeq v2, 300 cycle reagent Kit (Illumina) in a paired-end fashion (150 × 2).

### Whole Genome Amplification

Whole-genome sequencing of homogenized samples was performed as follows. Viral RNA was obtained as described above and then reverse transcribed and amplified using 2.5 µL of RNA as template in a 25 µL MS-RT-PCR reaction ^38^ (Superscript III high-fidelity RT-PCR kit, ThermoFisher). Primer sequences and concentrations were as follows: Uni12/Inf-1 5’-GGGGGGAGCAAAAGCAGG-3’ (0.06 µM); Uni12/Inf-3 5’-GGGGGGAGCGAAAGCAGG-3’ (0.14 µM); Uni13/Inf-1 5’-CGGGTTATTAGTAGAAACAAGG-3’ (0.2 µM). The cycling conditions were 55°C for 2 min, 42°C for 60 min, 94°C for 2 min, 5 cycles (94°C for 30 s, 44°C for 30 s, 68°C for 3.5 min), followed by 26 cycles (94°C for 30 s, 55°C for 30 s, 68°C for 3.5 min) and a final extension of 68°C for 10 min.

Amplicons were sequenced at either the Emory National Research Primate Center Genomics Core or at the Emory Integrated Genomics Core. Amplicon sequencing libraries were prepared using Nextera XT DNA library preparation kit (Illumina) according to the manufacturer’s protocol. NGS libraries were multiplexed and sequenced on a high-throughput Illumina NovaSeq 6000 (Illumina) in a paired-end 100 nucleotide run format.

### Analysis of barcode compositions

Sequences were processed using our custom software, BarcodeID, which is available in the GitHub repository (https://github.com/Lowen-Lab/BarcodeID). Briefly, BarcodeID uses BBTools to process fastq files. A custom Python script is then employed to screen and identify barcode sequences present in each sample, calculate diversity statistics, and generate summary tables. BBMerge screens reads for adapter sequences and merges forward and reverse reads using default settings, while BBDuk filters out reads with a low average quality (<30). BarcodeID subsequently verifies that the nucleotides at barcode and non-barcode sites match the expected nucleotides and possess sufficient quality (≥35 and ≥25, respectively). Reads with mismatches are excluded from the overall barcode sequence counts. However, BarcodeID collects all high-quality variant amplicons and calculates overall mismatch rates by site to identify if any mutants with non-barcode alleles are influencing observed barcode dynamics.

Genetic relationships of virus populations were evaluated by hierarchical clustering. A matrix of Euclidean distances of barcodes frequencies was used. The base R function *hclust* with method “complete” was used to perform hierarchical clustering. The R program “plot” was used to visualize the results. R version 4.1.3, (http://www.r-project.org).

### Whole genome variant analysis

Non-consensus variant analysis was conducted using a custom LoFreq-based pipeline ^39^. Adapter removal was performed using Cutadapt, followed by mapping the reads to the reference genome using BWA-MEM ^40^. Data formatting included using Samtools for fixing mate information, and LoFreq Viterbi for probabilistic realignment to correct mapping errors. The reads were then sorted with Samtools and indel qualities were inserted using LoFreq indelqual. Base and indel alignment qualities were added with LoFreq alnqual.

Variant calling was executed with LoFreq, generating a VCF file. Custom scripts filtered and formatted the VCF file to extract allele frequencies and coverage statistics. Only variants at a frequency of 0.05 with a coverage equal or above 400 were used. For detection of synonymous and nonsynonymous mutations we used the program SNPdat ^41^.

### Dissimilarity indexes

To establish genetic relationships between populations we used two dissimilarity indexes, Bray-Curtis for comparing barcode populations and Raup-Crick for whole genome pairwise comparison ^42,43^. The Raup-Crick dissimilarity index was chosen for whole genome data due to its robustness in calculating dissimilarity for unevenly sampled populations, ensuring that the calculated dissimilarities are accurate even when some samples have more or fewer available data points than others. Both indices consider the species shared between two populations. In the case of Bray-Curtis, the species were represented by unique barcode whereas in the case of Raup-Crick, species were represented by unique variants present in the genome, without taking into consideration those introduced to generate the barcodes.

Calculation of Bray-Curtis and Raup-Crick dissimilarity indexes was done using the vegan package version 2.6-4 in R.

### Plots

All figures were made using RStudio and the package ggplot2 (Wickham 2016) and aesthetically modified using Inkscape v.1.3.2 (https://inkscape.org).

### Data availability

The sequences are available through NCBI’s Short Read Archive (https://www.ncbi.nlm.nih.gov/sra) BioProject accession number PRJNAXXXX. All custom computer code necessary to reproduce the results presented in the manuscript are available on GitHub. The pipeline for the analysis of barcode composition is at (https://github.com/Lowen-Lab/BarcodeID). The pipeline for whole genome analysis is at (https://github.com/genferreri/Initial_LoFreq).

## References

1. Manicassamy, B., Manicassamy, S., Belicha-Villanueva, A., Pisanelli, G., Pulendran, B., and Garcia-Sastre, A. (2010). Analysis of in vivo dynamics of influenza virus infection in mice using a GFP reporter virus. Proc Natl Acad Sci U S A 107, 11531–11536. 10.1073/pnas.0914994107.

2. Karlsson, E.A., Meliopoulos, V.A., Savage, C., Livingston, B., Mehle, A., and Schultz-Cherry, S. (2015). Visualizing real-time influenza virus infection, transmission and protection in ferrets. Nat Commun 6, 6378. 10.1038/ncomms7378.

3. Gallagher, M.E., Brooke, C.B., Ke, R., and Koelle, K. (2018). Causes and Consequences of Spatial Within-Host Viral Spread. Viruses 10. 10.3390/v10110627.

4. Fukuyama, S., Katsura, H., Zhao, D., Ozawa, M., Ando, T., Shoemaker, J.E., Ishikawa, I., Yamada, S., Neumann, G., Watanabe, S., et al. (2015). Multi-spectral fluorescent reporter influenza viruses (Color-flu) as powerful tools for in vivo studies. Nat Commun 6, 6600. 10.1038/ncomms7600.

5. Zwart, M.P., and Elena, S.F. (2015). Matters of Size: Genetic Bottlenecks in Virus Infection and Their Potential Impact on Evolution. Annu Rev Virol 2, 161–179. 10.1146/annurev-virology-100114-055135.

6. Shi, Y.T., Harris, J.D., Martin, M.A., and Koelle, K. (2024). Transmission Bottleneck Size Estimation from De Novo Viral Genetic Variation. Mol Biol Evol 41. 10.1093/molbev/msad286.

7. Gutierrez, S., Michalakis, Y., and Blanc, S. (2012). Virus population bottlenecks during within-host progression and host-to-host transmission. Curr Opin Virol 2, 546–555. 10.1016/j.coviro.2012.08.001.

8. Takayama, I., Nguyen, B.G., Dao, C.X., Pham, T.T., Dang, T.Q., Truong, P.T., Do, T.V., Pham, T.T.P., Fujisaki, S., Odagiri, T., et al. (2021). Next-Generation Sequencing Analysis of the Within-Host Genetic Diversity of Influenza A(H1N1)pdm09 Viruses in the Upper and Lower Respiratory Tracts of Patients with Severe Influenza. mSphere 6. 10.1128/mSphere.01043-20.

9. Lauring, A.S. (2020). Within-Host Viral Diversity: A Window into Viral Evolution. Annu Rev Virol 7, 63–81. 10.1146/annurev-virology-010320-061642.

10. Farjo, M., Koelle, K., Martin, M.A., Gibson, L.L., Walden, K.K.O., Rendon, G., Fields, C.J., Alnaji, F.G., Gallagher, N., Luo, C.H., et al. (2024). Within-host evolutionary dynamics and tissue compartmentalization during acute SARS-CoV-2 infection. J Virol 98, e0161823. 10.1128/jvi.01618-23.

11. Ball, J.K., Holmes, E.C., Whitwell, H., and Desselberger, U. (1994). Genomic variation of human immunodeficiency virus type 1 (HIV-1): molecular analyses of HIV-1 in sequential blood samples and various organs obtained at autopsy. J Gen Virol 75 ( Pt 4), 67–79. 10.1099/0022-1317-75-4-867.

12. Excoiier, L., Foll, M., and Petit, R.J. (2009). Genetic Consequences of Range Expansions. Annu Rev Ecol Evol S 40, 481–501. 10.1146/annurev.ecolsys.39.110707.173414.

13. Fisher, R.A. (1937). The wave of advance of advantageous genes. Ann Eugenic 7, 355–369. DOI 10.1111/j.1469-1809.1937.tb02153.x.

14. Bialozyt, R., Ziegenhagen, B., and Petit, R.J. (2006). Contrasting eiects of long distance seed dispersal on genetic diversity during range expansion. J Evol Biol 19, 12–20. 10.1111/j.1420-9101.2005.00995.x.

15. Paulose, J., and Hallatschek, O. (2020). The impact of long-range dispersal on gene surfing. Proc Natl Acad Sci U S A 117, 7584–7593. 10.1073/pnas.1919485117.

16. Alpert, P., Bone, E., and Holzapfel, C. (2000). Invasiveness, invasibility and the role of environmental stress in the spread of non-native plants. Perspectives in Plant Ecology, Evolution and Systematics 3, 52–66. 10.1078/1433-8319-00004.

17. Lau, Y.H. (2008). Principles of population genetics, 4th edition. Econ Bot 62, 200–201.

18. Hallatschek, O., and Nelson, D.R. (2008). Gene surfing in expanding populations. Theor Popul Biol 73, 158–170. 10.1016/j.tpb.2007.08.008.

19. Ibrahim, K.M., Nichols, R.A., and Hewitt, G.M. (1996). Spatial patterns of genetic variation generated by diierent forms of dispersal during range expansion. Heredity 77, 282–291. DOI 10.1038/hdy.1996.142.

20. McCrone, J.T., Woods, R.J., Martin, E.T., Malosh, R.E., Monto, A.S., and Lauring, A.S. (2018). Stochastic processes constrain the within and between host evolution of influenza virus. Elife 7. ARTN e35962 10.7554/eLife.35962.

21. Varble, A., Albrecht, R.A., Backes, S., Crumiller, M., Bouvier, N.M., Sachs, D., Garcia-Sastre, A., and tenOever, B.R. (2014). Influenza A virus transmission bottlenecks are defined by infection route and recipient host. Cell Host Microbe 16, 691–700. 10.1016/j.chom.2014.09.020.

22. Fitzmeyer, E.A., Gallichotte, E.N., and Ebel, G.D. (2023). Scanning barcodes: A way to explore viral populations. PLoS Pathog 19, e1011291. 10.1371/journal.ppat.1011291.

23. Roder, A.E., Johnson, K.E.E., Knoll, M., Khalfan, M., Wang, B., Schultz-Cherry, S., Banakis, S., Kreitman, A., Mederos, C., Youn, J.H., et al. (2023). Optimized quantification of intra-host viral diversity in SARS-CoV-2 and influenza virus sequence data. mBio 14, e0104623. 10.1128/mbio.01046-23.

24. Lauck, M., Alvarado-Mora, M.V., Becker, E.A., Bhattacharya, D., Striker, R., Hughes, A.L., Carrilho, F.J., O’Connor, D.H., and Pinho, J.R. (2012). Analysis of hepatitis C virus intrahost diversity across the coding region by ultradeep pyrosequencing. J Virol 86, 3952–3960. 10.1128/JVI.06627-11.

25. Rogers, M.B., Song, T., Sebra, R., Greenbaum, B.D., Hamelin, M.E., Fitch, A., Twaddle, A., Cui, L., Holmes, E.C., Boivin, G., and Ghedin, E. (2015). Intrahost dynamics of antiviral resistance in influenza A virus reflect complex patterns of segment linkage, reassortment, and natural selection. mBio 6. 10.1128/mBio.02464-14.

26. Waters, J.M., Fraser, C.I., and Hewitt, G.M. (2013). Founder takes all: density-dependent processes structure biodiversity. Trends Ecol Evol 28, 78–85. 10.1016/j.tree.2012.08.024.

27. Lefèvre, F., Fady, B., Fallour-Rubio, D., Ghosn, D., and Bariteau, M. (2004). Impact of founder population, drift and selection on the genetic diversity of a recently translocated tree population. Heredity 93, 542–550. 10.1038/sj.hdy.6800549.

28. Amato, K.A., Haddock, L.A., 3rd, Braun, K.M., Meliopoulos, V., Livingston, B., Honce, R., Schaack, G.A., Boehm, E., Higgins, C.A., Barry, G.L., et al. (2022). Influenza A virus undergoes compartmentalized replication in vivo dominated by stochastic bottlenecks. Nat Commun 13, 3416. 10.1038/s41467-022-31147-0.

29. Ganti, K., Bagga, A., Carnaccini, S., Ferreri, L.M., Geiger, G., Joaquin Caceres, C., Seibert, B., Li, Y., Wang, L., Kwon, T., et al. (2022). Influenza A virus reassortment in mammals gives rise to genetically distinct within-host subpopulations. Nat Commun 13, 6846. 10.1038/s41467-022-34611-z.

30. Nichols, R.A., and Hewitt, G.M. (1994). The Genetic Consequences of Long-Distance Dispersal during Colonization. Heredity 72, 312–317. DOI 10.1038/hdy.1994.41.

31. Garant, D., Kruuk, L.E.B., Wilkin, T.A., McCleery, R.H., and Sheldon, B.C. (2005). Evolution driven by diierential dispersal within a wild bird population. Nature 433, 60–65. 10.1038/nature03051.

32. Malmberg, R., Simonsson, B., and Berglund, E. (1963). Airways obstruction and uneven gas distribution in the lung. Thorax 18, 168–171. 10.1136/thx.18.2.168.

33. Sims, A., Tornaletti, L.B., Jasim, S., Pirillo, C., Devlin, R., Hirst, J.C., Loney, C., Wojtus, J., Sloan, E., Thorley, L., et al. (2023). Superinfection exclusion creates spatially distinct influenza virus populations. PLoS Biol 21, e3001941. 10.1371/journal.pbio.3001941.

34. Sjaastad, L.E., Fay, E.J., Fiege, J.K., Macchietto, M.G., Stone, I.A., Markman, M.W., Shen, S., and Langlois, R.A. (2018). Distinct antiviral signatures revealed by the magnitude and round of influenza virus replication in vivo. Proc Natl Acad Sci U S A 115, 9610–9615. 10.1073/pnas.1807516115.

35. Belser, J.A., Barclay, W., Barr, I., Fouchier, R.A.M., Matsuyama, R., Nishiura, H., Peiris, M., Russell, C.J., Subbarao, K., Zhu, H., and Yen, H.L. (2018). Ferrets as Models for Influenza Virus Transmission Studies and Pandemic Risk Assessments. Emerg Infect Dis 24, 965–971. 10.3201/eid2406.172114.

36. Belser, J.A., Eckert, A.M., Huynh, T., Gary, J.M., Ritter, J.M., Tumpey, T.M., and Maines, T.R. (2020). A Guide for the Use of the Ferret Model for Influenza Virus Infection. Am J Pathol 190, 11–24. 10.1016/j.ajpath.2019.09.017.

37. Perez, D.R., Seibert, B., Ferreri, L., Lee, C.W., and Rajao, D. (2020). Plasmid-Based Reverse Genetics of Influenza A Virus. Methods Mol Biol 2123, 37–59. 10.1007/978-1-0716-0346-8_4.

38. Zhou, B., and Wentworth, D.E. (2012). Influenza A virus molecular virology techniques. Methods Mol Biol 865, 175–192. 10.1007/978-1-61779-621-0_11.

39. Wilm, A., Aw, P.P., Bertrand, D., Yeo, G.H., Ong, S.H., Wong, C.H., Khor, C.C., Petric, R., Hibberd, M.L., and Nagarajan, N. (2012). LoFreq: a sequence-quality aware, ultra-sensitive variant caller for uncovering cell-population heterogeneity from high-throughput sequencing datasets. Nucleic Acids Res 40, 11189–11201. 10.1093/nar/gks918.

40. Li, H., and Durbin, R. (2009). Fast and accurate short read alignment with Burrows-Wheeler transform. Bioinformatics 25, 1754–1760. 10.1093/bioinformatics/btp324.

41. Doran, A.G., and Creevey, C.J. (2013). Snpdat: easy and rapid annotation of results from de novo snp discovery projects for model and non-model organisms. BMC Bioinformatics 14, 45. 10.1186/1471-2105-14-45.

42. Bray, J.R., and Curtis, J.T. (1957). An Ordination of the Upland Forest Communities of Southern Wisconsin. Ecol Monogr 27, 326–349. 10.2307/1942268.

43. Raup, D.M., and Crick, R.E. (1979). Measurement of Faunal Similarity in Paleontology. J Paleontol 53, 1213–1227.

