## Supplementary material for "Dispersal of influenza virus populations within the respiratory tract shapes their evolutionary potential": Figure S1, Figure S2, Figure S3, Figure S4, Figure S5, Figure S6, Figure S7

**Supplementary Figures**

 
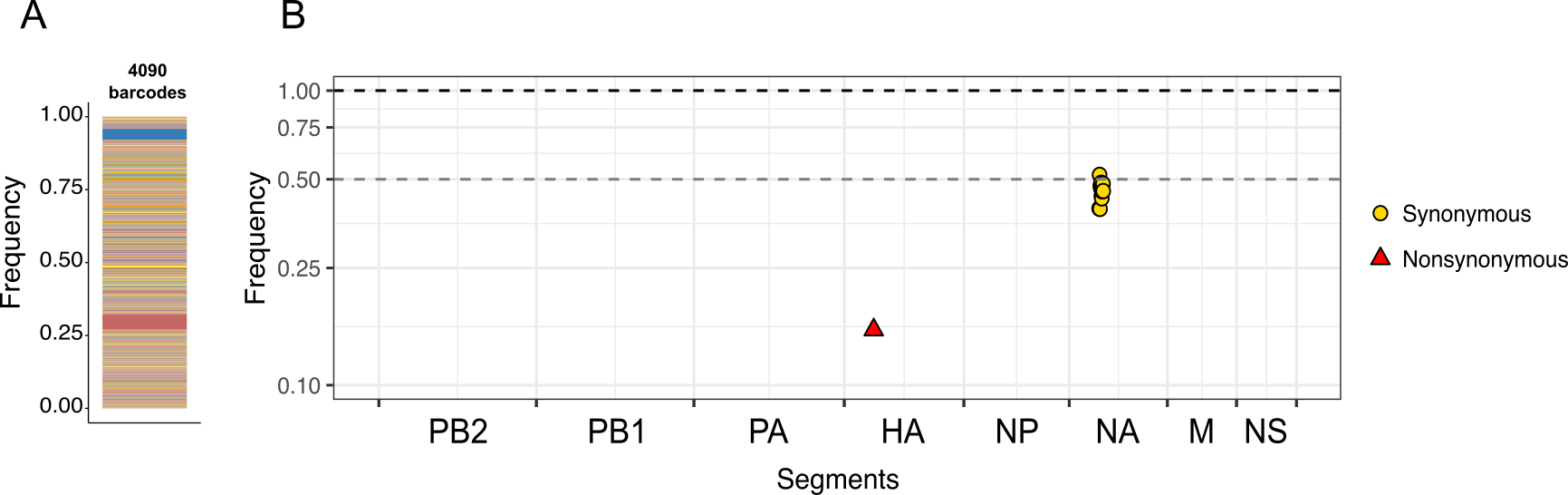

**Figure S1. Characterizing genotypes in the inoculum.** A. Barcodes detected in the viral inoculum. Lines of different colors represent unique barcodes. Due to the high count of barcodes, colors were recycled. The thickness of lines represents the frequency of each barcode. B. Variants detected above 0.05 frequency in the inoculum throughout the viral genome. Dashed grey line indicates a frequency of 0.5. Black dashed line indicates a frequency of 1.0.

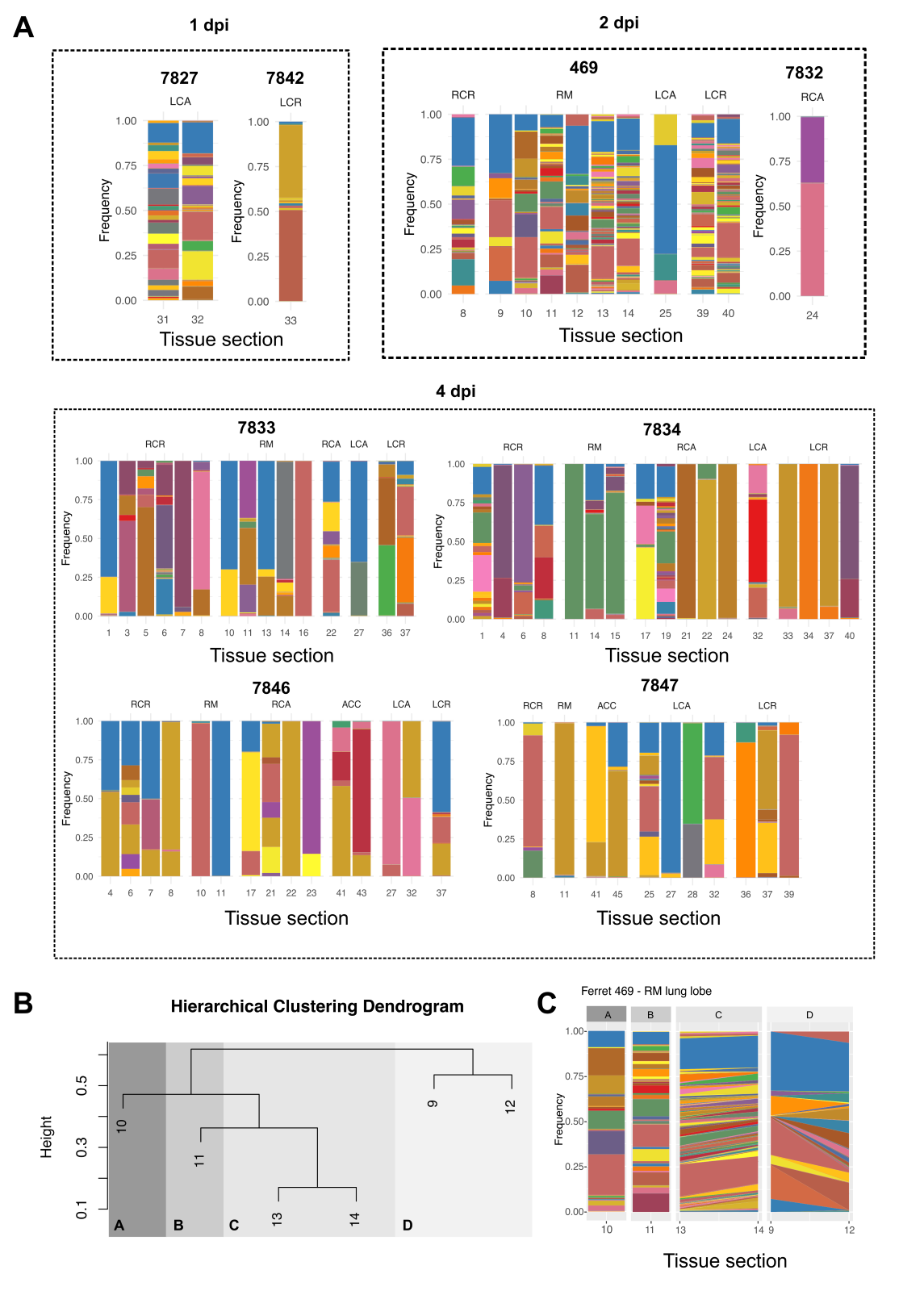

**Figure S2. Barcode composition in the lung of intranasally inoculated ferrets.** (A) Barcode detection in lung sections of individual ferrets. (B) Relationships among viral populations sampled from the right middle lobe of ferret 469, as assessed by hierarchical clustering of barcode compositions. Height shows distance between clusters. Subsections A-D show that different populations identified. (C) Continuity of barcodes within clusters identified in (B). Hierarchical clustering was applied where at least three sections per lobe were positive and only to animals sampled at 1 or 2 dpi since stochastic loss of barcodes by 4 dpi prevents detection of genetic relatedness.

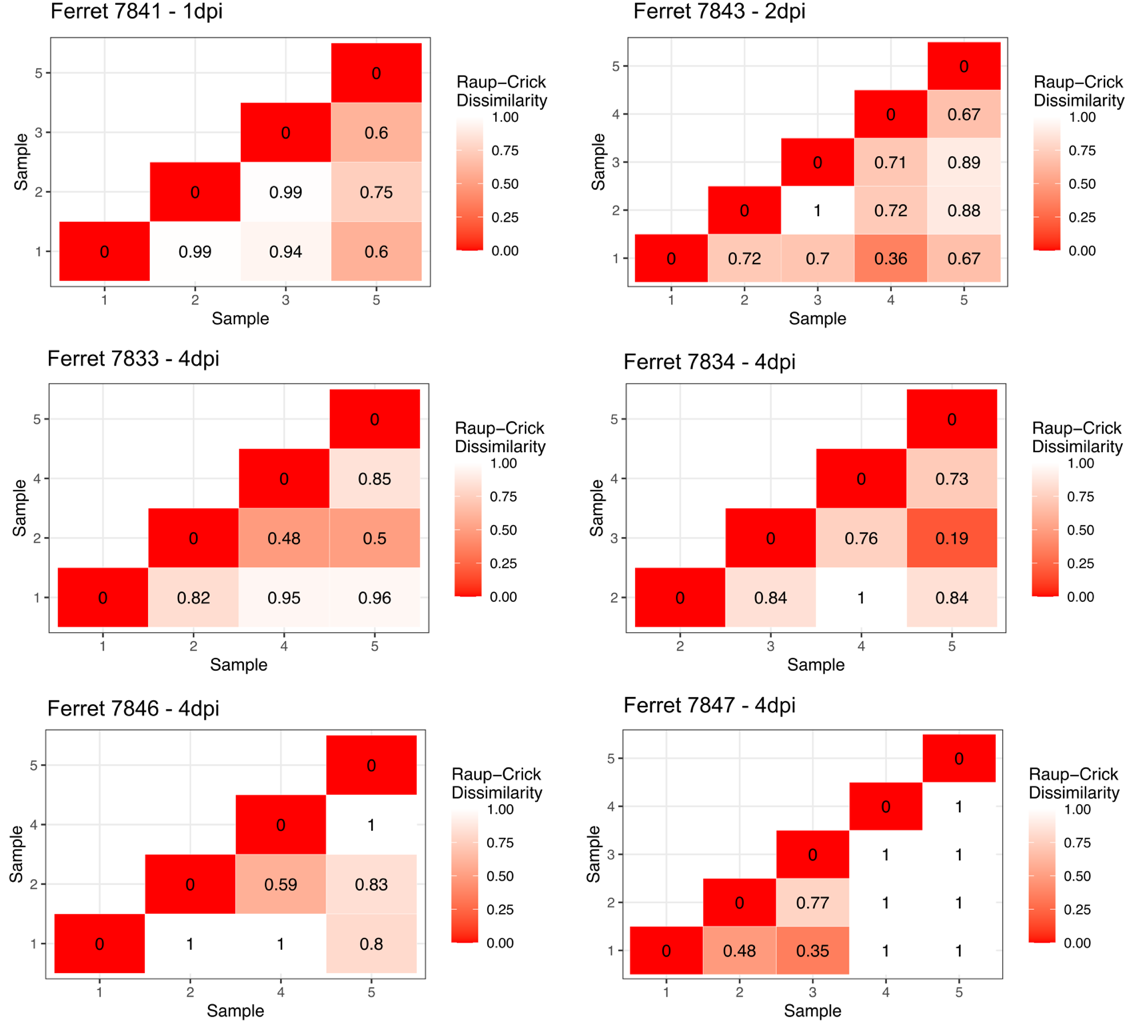

**Figure S3. Heterogeneous subpopulations are formed as the result of de novo mutations in trachea.** Raup-Crick dissimilarity index for the analysis of virus populations in trachea. De novo variants in the HA, NP, NA, M and NS gene segments were considered.

 
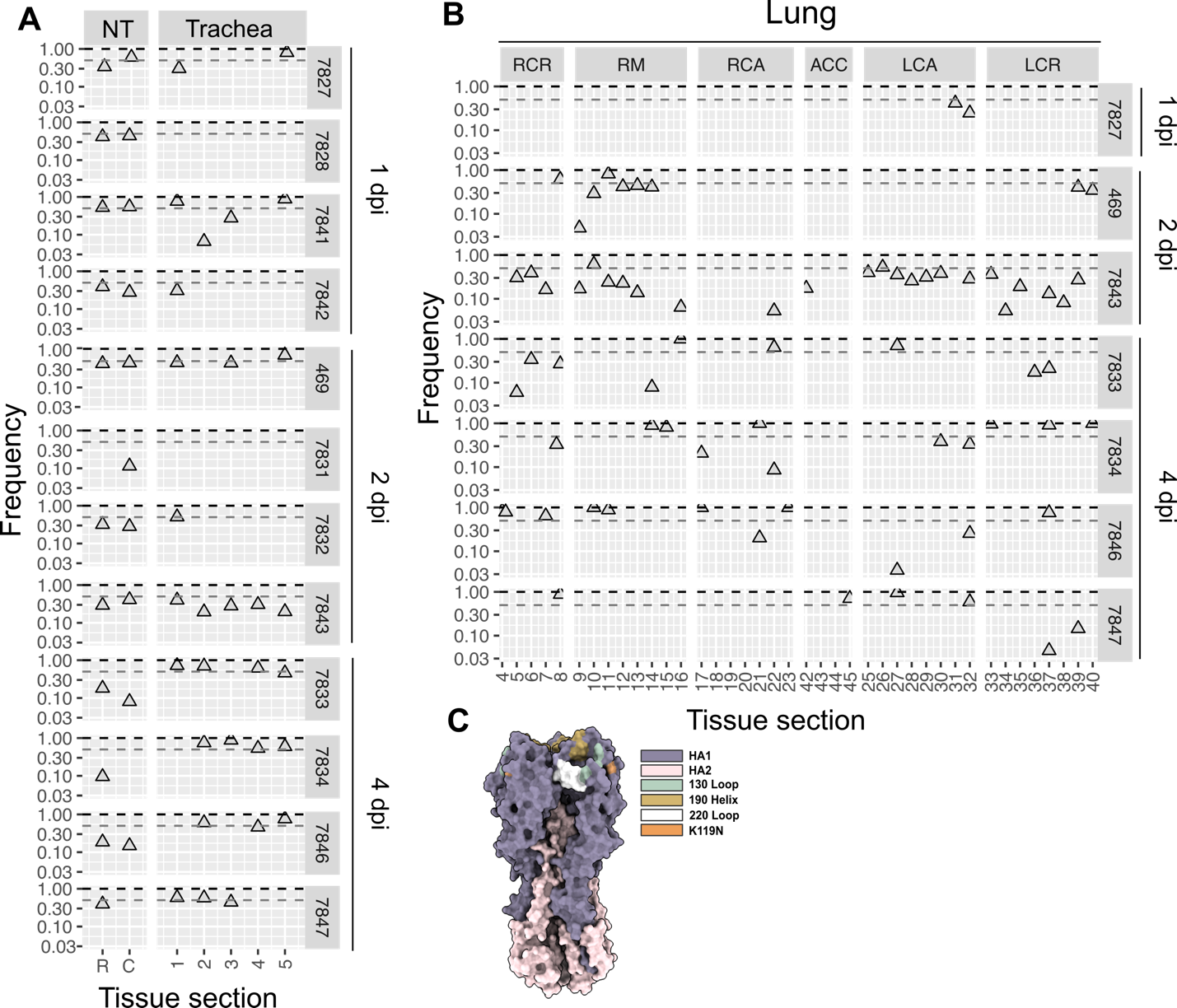

**Figure S4. Detection of mutation HA K119N across the respiratory tract.** (A) Frequency of mutation HA K119N in the nasal turbinates (NT) and trachea. (B) Frequency of mutation HA K119N in the lungs. RCR = right cranial; RM = right middle; RCA = right caudal; ACC = accessory; LCA = left caudal; LCR = left cranial. (C) Location of HA K119N in the HA trimer. The receptor binding site is delimited by the 130 loop, 190 helix and the 220 loop.

 
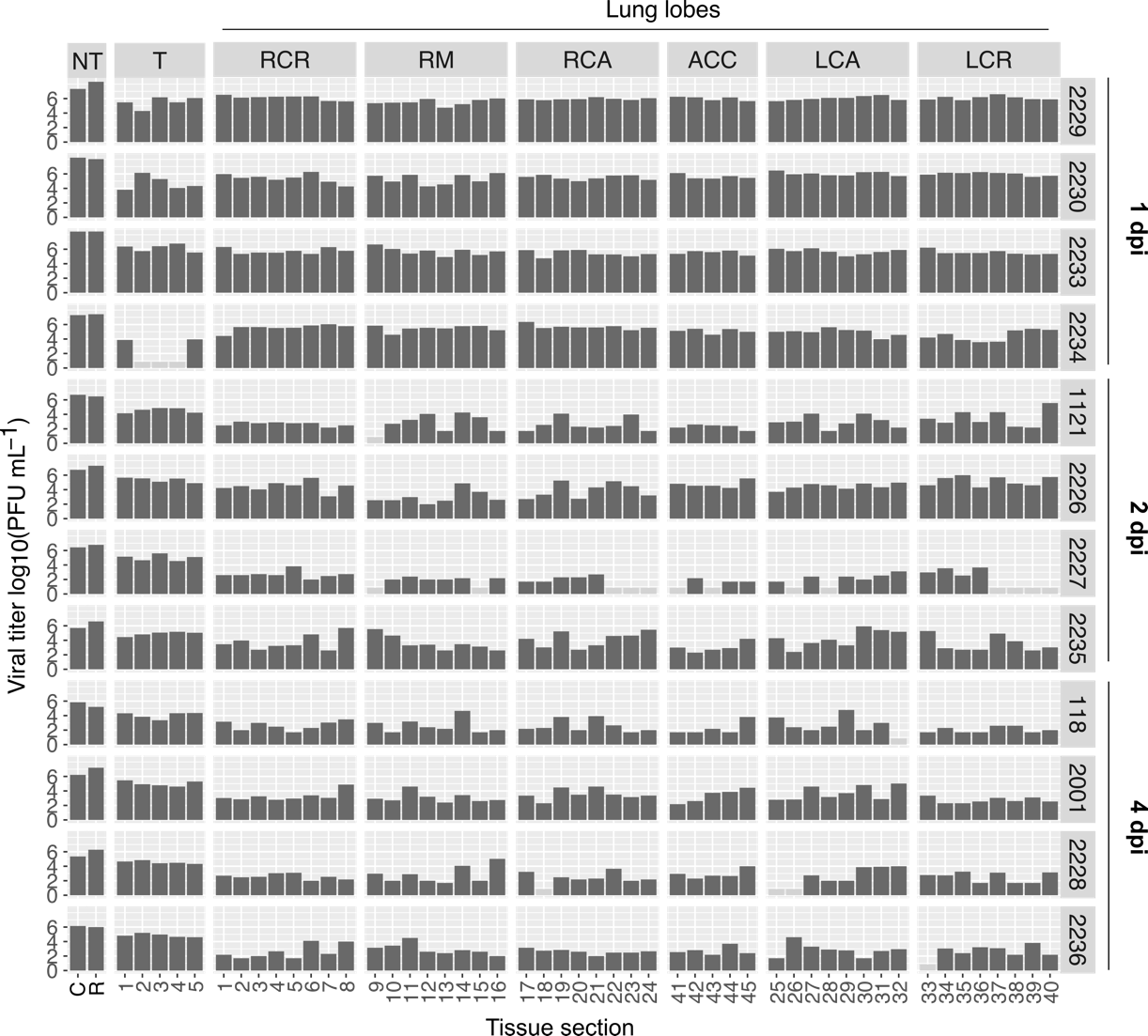

**Figure S5. Extensive infection after aerosol inoculation.** Dark grey bars depict positive samples. Eight sections were sampled from each lung lobe, except the accessory lobe, from which five sections were collected. NT = nasal turbinates; T = trachea. Lobes are designated as: RCR = right cranial; RM = right middle; RCA = right caudal; ACC = accessory; LCA = left caudal; LCR = left cranial. Numbers on the secondary y-axis show ferret identification numbers. Time of sampling is shown at the extreme right (dpi = days post-inoculation).

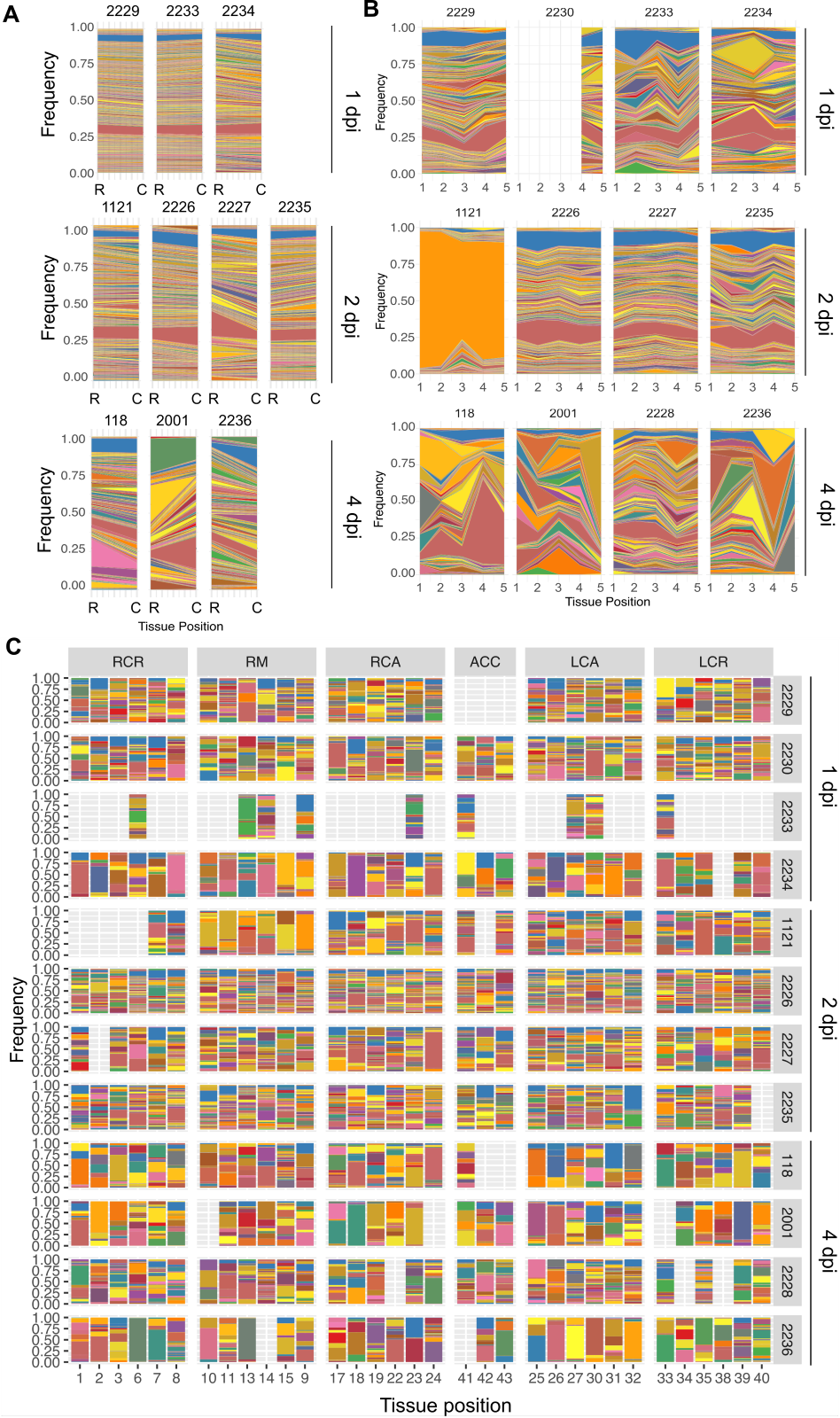

**Figure S6. Aerosol inoculation seeds largely uniform populations in the nasal turbinate and trachea but heterogeneous populations throughout the lower respiratory tract.** Barcode composition in the nasal turbinates (A), in trachea (B) and lungs (C). In the nasal turbinates R = rostral and C = caudal. The right y-axis displays the ferret ID numbers. RCR = right cranial; RM = right middle; RCA = right caudal; ACC = accessory; LCA = left caudal; LCR = left cranial.

 
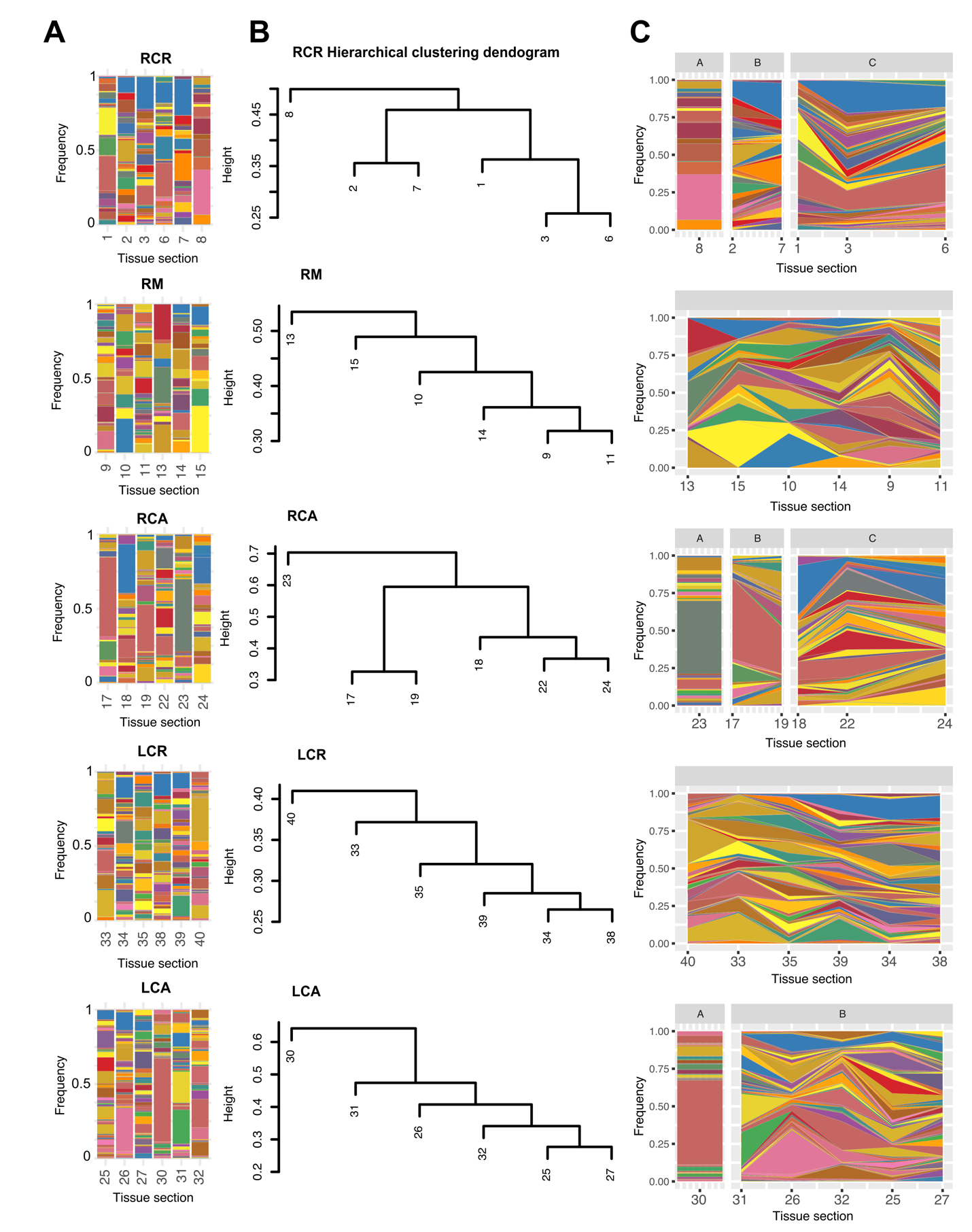

**Figure S7. Aerosol inoculation seeds heterogeneous populations in the lungs.** (A) Barcode composition by tissue section for the lung lobes LCA and LCR of ferret 2230, which was sampled at 1 dpi. (B) Distance matrices of barcode frequencies were used to evaluate genetic relationships by hierarchical clustering. Height shows distance between clusters. (C) Stacked plots organized according to the genetic relationships identified by hierarchical clustering for all lobes except the accessory lobe (ACC).
